# Aberrant DNA methylation of miRNAs in Fuchs endothelial corneal dystrophy

**DOI:** 10.1101/638486

**Authors:** Peipei Pan, Daniel J. Weisenberger, Siyu Zheng, Marie Wolf, David G. Hwang, Jennifer R. Rose-Nussbaumer, Ula V. Jurkunas, Matilda F. Chan

## Abstract

Homeostatic maintenance of corneal endothelial cells is essential for maintenance of corneal deturgescence and corneal transparency. In Fuchs endothelial corneal dystrophy (FECD), an accelerated loss and dysfunction of endothelial cells leads to progressively severe visual impairment. An abnormal accumulation of extracellular matrix is a distinctive hallmark of the disease, however the molecular pathogenic mechanisms underlying this phenomenon are not fully understood. We recently reported characteristic patterns of DNA methylation changes in the corneal endothelial cells of patients with FECD. Here, we investigate genome-wide and sequence-specific DNA methylation changes of miRNA genes in corneal endothelial samples derived from patients with FECD. We show that the majority of miRNA genes are hypermethylated at their promoter regions in FECD. More specifically, *miR-199B* is an extensively hypermethylated miRNA gene at its promoter region and its mature transcript miR-199b-5p was previously found to be almost completely silenced in FECD. Using a cell-based assay, we find that miR-199b-5p directly inhibits the expression of two epithelial mesenchymal transition (EMT)-inducing genes, *Snai1* and *ZEB1*. Taken together, these findings suggest a novel regulatory mechanism of matrix protein production by corneal endothelial cells in which miR-199b-5p hypermethylation leads to its down-regulated expression and thereby the decreased expression of miR-199b-5p target genes, including *Snai1* and *ZEB1*. Our results support miR-199b-5p as a potential therapeutic target to prevent or slow down the progression of FECD disease.

**Author summary:** Fuchs endothelial corneal dystrophy (FECD) due to corneal endothelial cell degeneration is one of the most common heritable causes of corneal visual loss and a leading indication for corneal transplantation. The progressive loss of corneal endothelial cells is accompanied by an abnormal deposition of extracellular matrix in the form of guttae. Here we discover that miRNA gene promoters are frequent targets of aberrant DNA methylation in FECD. In particular, we describe a novel epigenetic mechanism used by corneal endothelial cells to regulate extracellular matrix production. We find that miRNA-199b-5p functions as a negative regulator of Snai1 and ZEB1, two zinc finger transcription factors that have been shown to lead to increased production of extracellular matrix proteins. Furthermore, *miR-199B* was extensively hypermethylated in FECD and its mature transcript miR-199b-5p directly binds to the 3′-UTRs of *Snai1* and *ZEB1* genes. Ultimately, this may negatively modulate Snai1- and ZEB1-mediated production of extracellular matrix proteins. This work is the first to identify an important role of DNA methylation in the epigenetic regulation of miRNA-target genes in FECD and to describe a potential epigenetic biomarker for the treatment of FECD patients.

## Introduction

Corneal transparency is critical for good visual acuity. The corneal endothelium regulates the hydration status of the cornea and has an essential role in maintaining corneal deturgescence and preventing edema that can degrade corneal transparency. It is the innermost layer of the cornea and is composed of a single layer of cells that pump excess fluid out of the cornea through active ion-transport processes [1, 2].

Fuchs endothelial corneal dystrophy (FECD) is a bilateral, slowly progressive disorder in which the corneal endothelial cells are diseased and become less efficient at removing fluid. As a result, the highly ordered arrangement of collagen fibers in the corneal stromal layer become disrupted, leading to corneal opacification and vision loss [3]. Other clinical phenotypic changes that occur in FECD include an excessive accumulation of extracellular matrix (ECM), formation of central excrescences (corneal guttae), thickening of Descemet’s membrane, and corneal scarring [4]. At earlier stages of FECD, the formation of corneal guttae can cause light scatter and optical aberrations that can impair vision, even in the absence of overt corneal edema. In later FECD, overt endothelial dysfunction and resultant corneal edema contribute significantly to visual loss. Corneal endothelial cells are largely non-regenerative *in vivo* and their loss is often irreversible [5]. Medical management is often inadequate and corneal endothelial transplantation remains the main therapeutic option to restore vision in patients with advanced FECD. FECD is a leading indication for corneal transplantation in the United States [6].

FECD is a multi-factorial disease that is associated with a variety of reported spontaneous and inherited mutations [7, 8] and can manifest as both early- and late-onset forms [9]. Mutations in the more common late-onset FECD have been identified in several genes including *SLC4A11* [10, 11], *ZEB1* (also named as *TCF8*) [12], *LOXHD1* [13], and *AGBL1* [14]. An expanded trinucleotide repeat in the third intron of transcription factor 4 (*TCF4*, also referred to as E2-2) has also been found to be strongly associated with many cases of late-onset FECD [15, 16]. In addition to genetic variations, environmental factors such as oxidative stress have also been implicated in the pathogenesis of FECD through complex cellular and biochemical responses [17–20]. The external location of the cornea renders it directly exposed to the environment and thus more susceptible to external stimuli.

DNA methylation is an epigenetic change that facilitates cellular adaptation to changing environments and has repeatedly been linked to human diseases and aging [21–23]. This has raised interest in understanding the potential contribution of DNA methylation to the development of late-onset eye diseases such as glaucoma [24, 25], age-related macular degeneration [26–28], and cataract [29]. Epigenetic factors may help explain the phenotypic variation seen amongst cohorts with identical genotypes, including in FECD. We previously identified DNA methylation changes that occur in the corneal endothelial tissue of patients with late-onset FECD using a genome-wide DNA methylation array [30]. Furthermore, many of these changes occurred in microRNA (miRNA) sequences [30].

Multiple studies over the past decade have demonstrated that miRNAs profoundly influence the cellular responses of tissues to physiologic and pathophysiologic stresses in multiple disease states [31, 32]. Stress-dependent regulation can involve upregulation or downregulation of miRNA expression and lead to downstream signaling effects on mRNA targets [31]. Accumulating evidence from human and animal studies have shown that DNA methylation-associated silencing of miRNAs contributes to disease pathogenesis [33]. In the present study, we further analyzed the subset of miRNA data to test the hypothesis that aberrant DNA methylation of miRNAs contributes to FECD pathogenesis. We found that the majority of differentially methylated miRNAs display promoter DNA hypermethylation in FECD. Moreover, we identified *miR-199B* to be extensively hypermethylated in FECD. Using *in silico* and functional assays, we determined that miR-199b-5p directly targets and negatively regulates the expression of two key transcription factors that control ECM production in FECD. Taken together, these findings demonstrate a novel mechanism of epigenetic regulation of ECM production in FECD pathogenesis and identifies miR-199b-5p as a potential clinical biomarker of disease.

## Results

### Global DNA methylation patterns of miRNA sequences are altered in FECD

Our prior genome-scale analysis of the DNA methylation landscape of corneal endothelial tissue found a significant difference between FECD and normal control patients [30]. In particular, we identified a high number of differentially methylated miRNA sequences [30]. Because DNA methylation plays a central role in regulating miRNA expression [34, 35] and widespread miRNA downregulation has been observed in FECD [36], we performed a subanalysis of the Illumina Infinium HumanMethylation450 (HM450) array data to focus on the 2,227 miRNA probes (targeting 463 miRNA genes in total). Sample pairwise correlation and hierarchical clustering analyses revealed differential genome-wide miRNA DNA methylation patterns in FECD samples compared with normal control samples (Fig 1). Further nonparametric principle component analyses showed that this variance was not attributable to the clinical variables of age, sex, pachymetry, or guttata grading (data not shown).

**Fig. 1.**
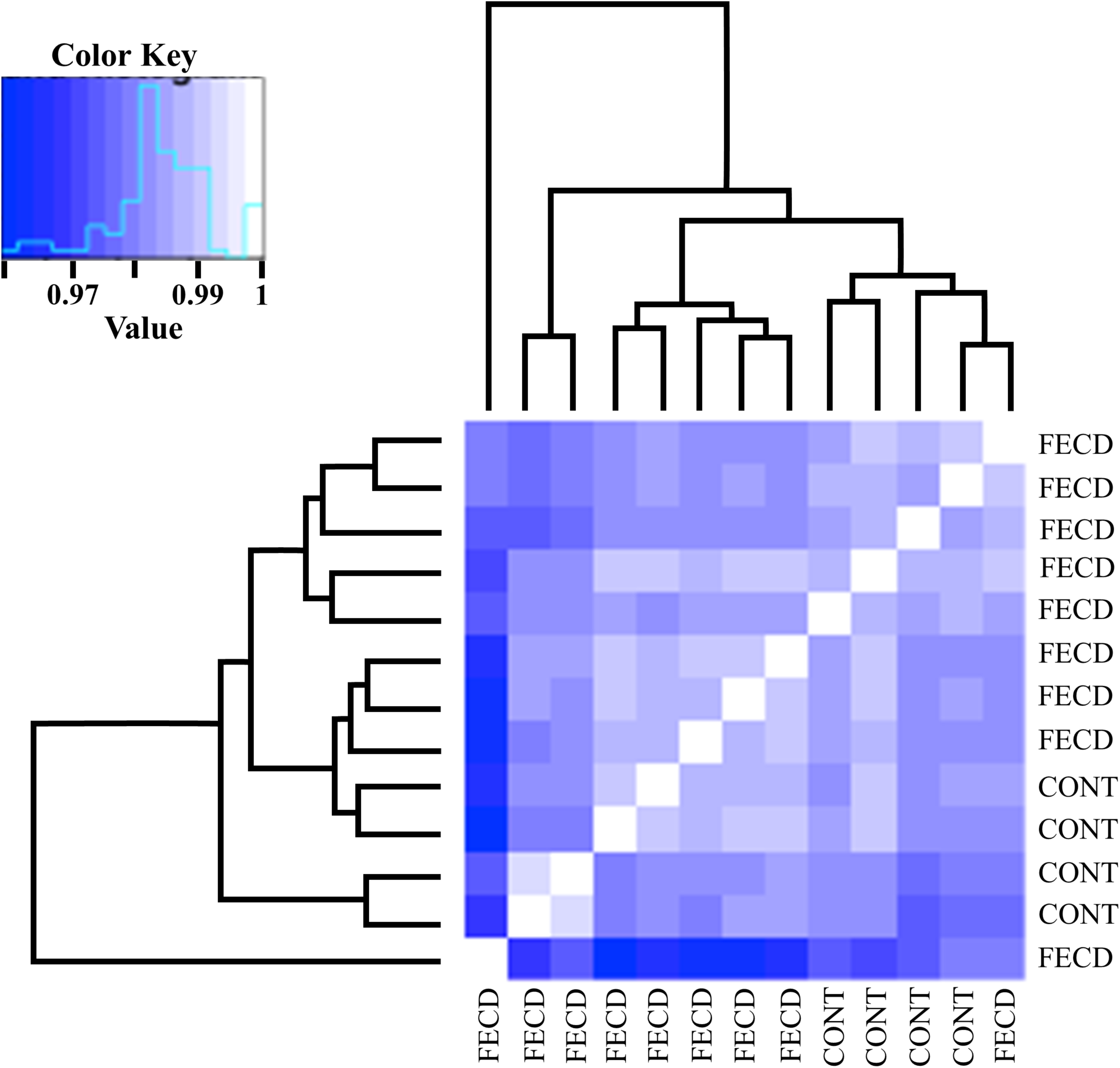
Heatmap visualization of pairwise Kendall rank correlation coefficients comparing miRNA gene methylation pattern between FECD and control corneal endothelium samples with dendrogram to show clustering (Euclidean distance). Control and FECD samples separate according to disease variable, with the exception of one FECD sample.

### The majority of differentially methylated miRNAs display promoter DNA hypermethylation in FECD

We next examined and compared the DNA methylation levels of individual miRNA sequences between FECD cases and controls by comparing single HM450 probes. Of the 2,227 miRNA-associated probes (targeting 463 miRNA genes), 216 probes (targeting 156 miRNA genes) were differentially methylated in the FECD samples (*P* < 0.05; Fig 2A). Of the 216 probes, the large majority (154 probes; 71%) were hypermethylated in the FECD samples, and a small minority (62 probes; 29%) of sequences were hypomethylated (Fig 2B). Almost all of the differentially methylated probes (208 probes; 96%) targeted miRNA promoter sequences (148/154 hypermethylated probes and 60/62 hypomethylated probes; Fig 2B). None of the miRNA probes were significantly differentially methylated with respect to age or sex, suggesting that these parameters are not major drivers of DNA methylation changes in miRNA genes in FECD patients (data not shown). Table 1 shows details on the 20 top ranking differentially methylated miRNA genes in the FECD samples as compared to controls. *miR199-B* was found to be the most hypermethylated miRNA gene and *miR-1182* was the most hypomethylated miRNA gene in FECD samples (Table 1).

**Fig 2.**
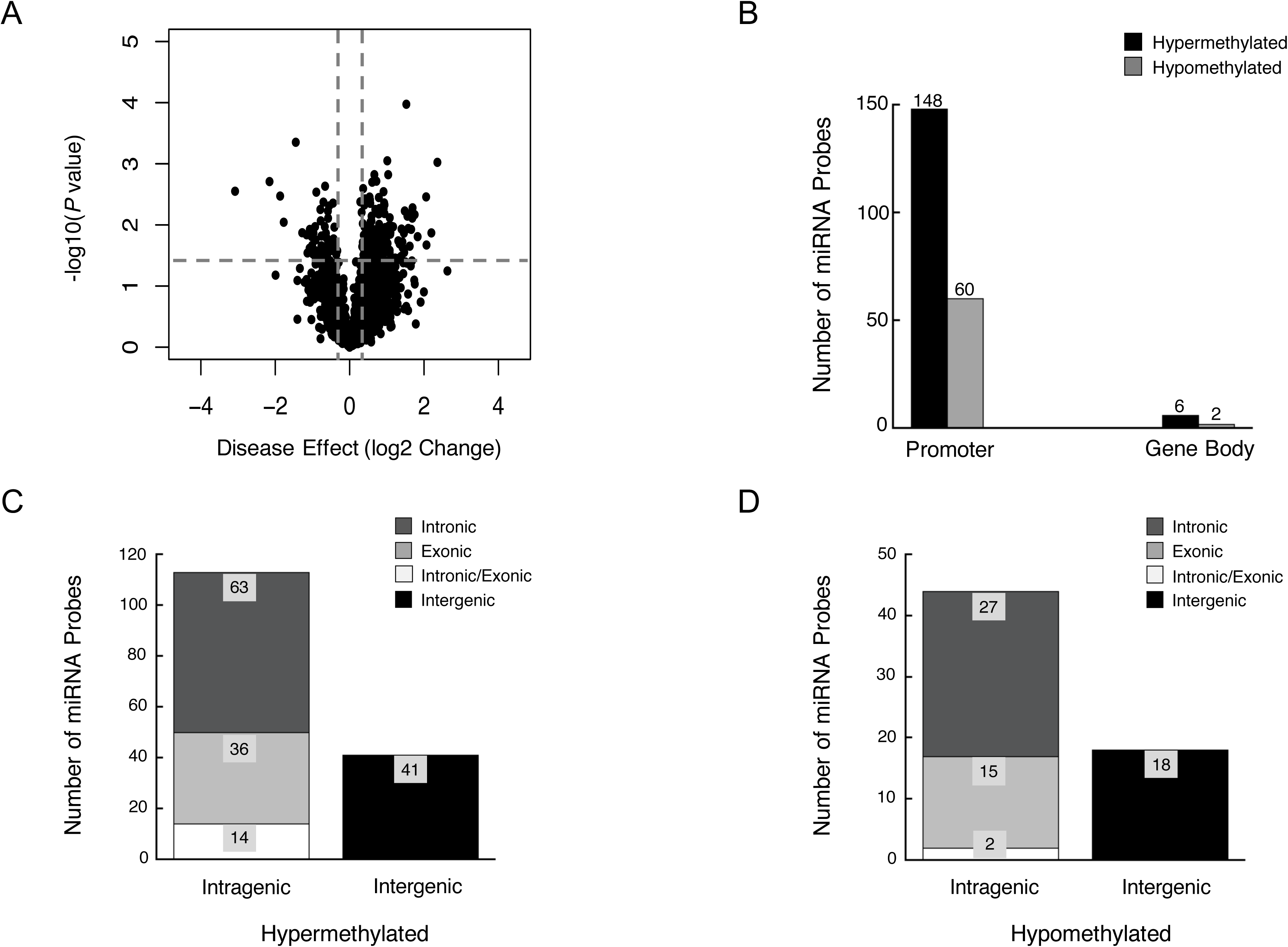
miRNA gene promoters are preferential target sites of aberrant DNA methylation in FECD. **(A)** Volcano plot shows the function of disease effect (log2 (fold change), x-axis) versus the statistical significance of the result (−log 10 (p-value), y-axis) for a total number of 2,227 probes targeting miRNAs genes on the array. Each dot represents an individual probe. Vertical dotted lines represent fold changes of ± 1.3, respectively. The horizontal dotted line indicates the p-value cutoff point (0.05; −log10(0.05) = 1.30103). The dots represent 216 selected differentially expressed probes with p-value < 0.05 and |fold-change| >1.3. **(B)** The blue dotted line indicate the p-value cutoff point (0.05; −log10(0.05) = 1.30103). Number of differentially methylated probes between FECD and control samples (p-value < 0.05), grouped by the probe targeting region in related miRNA genes. **(C)** Genome location of hypermethylated miRNA probes relative to the corresponding host genes. **(D)** Genomic location of hypomethylated miRNA probes relative to the corresponding host genes.

**Table 1.** Top 20 most significantly differently methylated miRNAs in FECD, with 18 of them being hypermethylated and 2 being hypomethylated.

| Gene | ProbeID | Host Genes | Fuchs_Coeff<sup>a</sup> | Fuchs_Pval | Fuchs_Qval |
| --- | --- | --- | --- | --- | --- |
| miR-199B | cg13718827 | DNM1 (antisense; intronic) | 2.3595 | *** | 0.00128 |
| miR-33B | cg04805065 | SREBF1 (sense; intronic) | 2.1974 | * | 0.12825 |
| miR-874 | cg18251187 | KLHL3 (sense; intronic) | 2.0571 | ** | 0.00944 |
| miR-1286 | cg08221669 | RTN4R (sense; intronic) | 1.7488 | ** | 0.02761 |
| miR-1306 | cg20689730 | DGCR8 (sense; exonic) | 1.6852 | ** | 0.03932 |
| miR-130A | cg10512089 | intergenic | 1.6548 | * | 0.09226 |
| miR-320B1 | cg25023761 | intergenic | 1.5556 | ** | 0.02459 |
| miR-641 | cg26620021 | AKT2 (sense; intronic) | 1.5535 | ** | 0.03169 |
| miR-942 | cg19582647 | TTF2 (sense; intronic) | 1.5362 | * | 0.82808 |
| miR-499 | cg11231913 | MYH7B (sense; intronic) | 1.5245 | *** | 0.00004 |
| miR-30B | cg25964744 | intergenic | 1.4986 | * | 0.75198 |
| miR-184 | cg23721598 | ANKRD34C-AS1(antisense; intronic) | 1.4823 | ** | 0.02005 |
| miR-194-2 | cg24803202 | MIR194-2HG (sense; exonic) | 1.4674 | * | 0.42089 |
| miR-25 | cg22638766 | MCM7 (sense; intronic) | 1.4593 | * | 0.13411 |
| miR-662 | cg26775123 | MSLNL (antisense; exonic) | 1.4426 | * | 0.80350 |
| miR-1471 | cg06046580 | intergenic | 1.3949 | * | 0.08720 |
| miR-199A1 | cg18544365 | DNM2 (antisense; intronic) | 1.3434 | * | 0.26307 |
| miR-320D1 | cg23483562 | intergenic | 1.2870 | * | 0.60071 |
| miR-1182 | cg20821842 | FAM89A (sense; exonic) | -3.0805 | ** | 0.00667 |
| miR-193A | cg08667128 | intergenic | -1.4483 | *** | 0.00030 |

<sup>a</sup>\* $P \leq 0.05$
\*\* $P \leq 0.01$
\*\*\* $P \leq 0.001$

### Genomic locations of differentially methylated miRNA genes with respect to their host genes

The majority of miRNAs are located within intronic or exonic regions of protein-coding genes (host genes), and increasing evidence suggests a functional relationship between miRNAs and their host genes [37]. Therefore, we next sought to examine the spatial relationship of the differentially methylated miRNA probes with their host genes. We mapped all 216 differentially methylated miRNA probes to their host genes using the Ensembl Genome Browser (https://uswest.ensembl.org/index.html). Of the 154 hypermethylated miRNA probes, 74% (113 probes) corresponded to intragenic sequences and 27% (41 probes) occurred in intergenic sequences (Fig 2C). Of the intragenic probes, 41% (63 probes) occurred within intronic sequences, 23% (36 probes) within exonic sequences, and 9% (14 probes) in intron/exon boundaries (Fig 2C). Similar to the hypermethylated probes, the majority of hypomethylated miRNA probes occurred in intragenic sequences (71%, 44 out of 62 probes) and intronic sequences (44% intronic, 24% exonic, 3% in intron/exon boundary sequences) (Fig 2D). Together, these data show that the majority of differentially methylated miRNA probes occur in intragenic and intronic sequences of their host genes. Our prior genome-scale analysis of the DNA methylation landscape of corneal endothelial tissue found a significant difference between FECD and normal control patients [30].

### Gene body regions of miRNA host genes are frequent targets of aberrant DNA methylation in FECD

Emerging evidence has revealed mutually regulatory roles between particular miRNAs and their host genes [38–41]. Therefore, to decipher the function of DNA methylation in the epigenetic regulation of miRNAs and their host genes, we next assessed the methylation status of host genes for the 156 differentially methylation miRNA genes in FECD. The list of 156 host genes was curated using publicly available miRNA databases, including Ensembl Genome Browser (https://uswest.ensembl.org/index.html), miRIAD (http://www.miriad-database.org), and miRStart (http://mirstart.mbc.nctu.edu.tw/about.php). A total of 1,823 probes mapped to CpG sites in the set of 156 host genes. A volcano plot display of the DNA methylation status of the 1,823 probes showed that 239 probes were differentially methylated in FECD samples compared to control samples (*P* < 0.05; Fig 3A). In fact, most (188; 79%) of the 239 differentially methylated CpG sites were hypermethylated in FECD samples, with only a minority (51; 21%) of them hypomethylated. Alignment of the probe sequences to the human genome database revealed that the vast majority (192; 80%) of the 239 differentially methylated CpG sites were located in the gene body regions of miRNA host genes (Fig 3B). A substantial proportion (161; 86%) of the 188 hypermethylated probes targeted CpG sites within gene bodies, whereas only 14% of them were mapped to the promoter regions of corresponding miRNA host genes. Similarly, a high percentage (31; 61%) of the 51 hypomethylated probes also mapped to gene body sequences of miRNA host genes. Table 2 provides detailed information on the top 20 differentially methylated miRNA host genes identified in the FECD samples as compared to controls. Given that miRNAs may be co-regulated with their host genes, we further analyzed the DNA methylation data for co-methylation patterns. We identified a subset of miRNAs and their host genes to be co-methylated at their corresponding promoter CpG sites, suggesting that DNA methylation may play an important role in regulating the co-expression of miRNAs and their host genes in FECD (Fig 3C).

**Fig 3.**
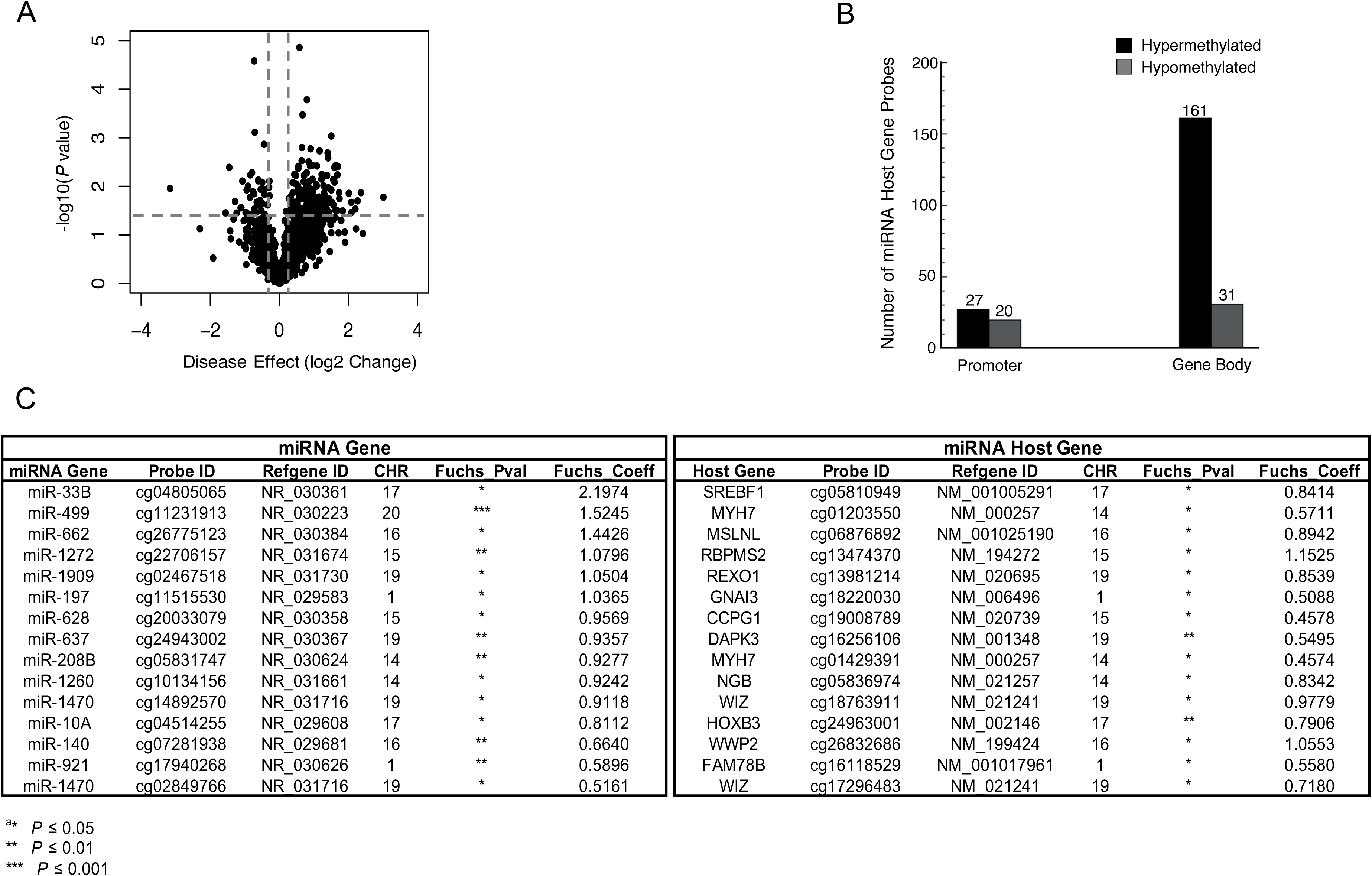
MiRNAs and host genes were co-methylated at corresponding promoter regions in FECD. **(A)** Volcano plot showing the function of disease effect (log2 (fold change), x-axis) versus the statistical significance of the result (−log 10 (p-value), y-axis) for a total number of 1,823 probes targeting miRNAs host genes on the array. Each dot represents a probe. Vertical dotted lines represent fold changes of ± 1.3, respectively. The horizontal dotted line indicate the p-value cutoff point (0.05; −log10(0.05) = 1.30103). The dots represent 239 selected differentially expressed miRNA host genes with p-value < 0.05 and |fold-change| >1.3. **(B)** Number of differentially methylated probes associated with miRNA host genes between FECD and control samples (*P* < 0.05), grouped by the probe targeting region in related miRNA host genes. **(C)** A subset of 15 miRNAs and their host genes with co-methylation at promoter CpG sites in FECD.

**Table 2.**
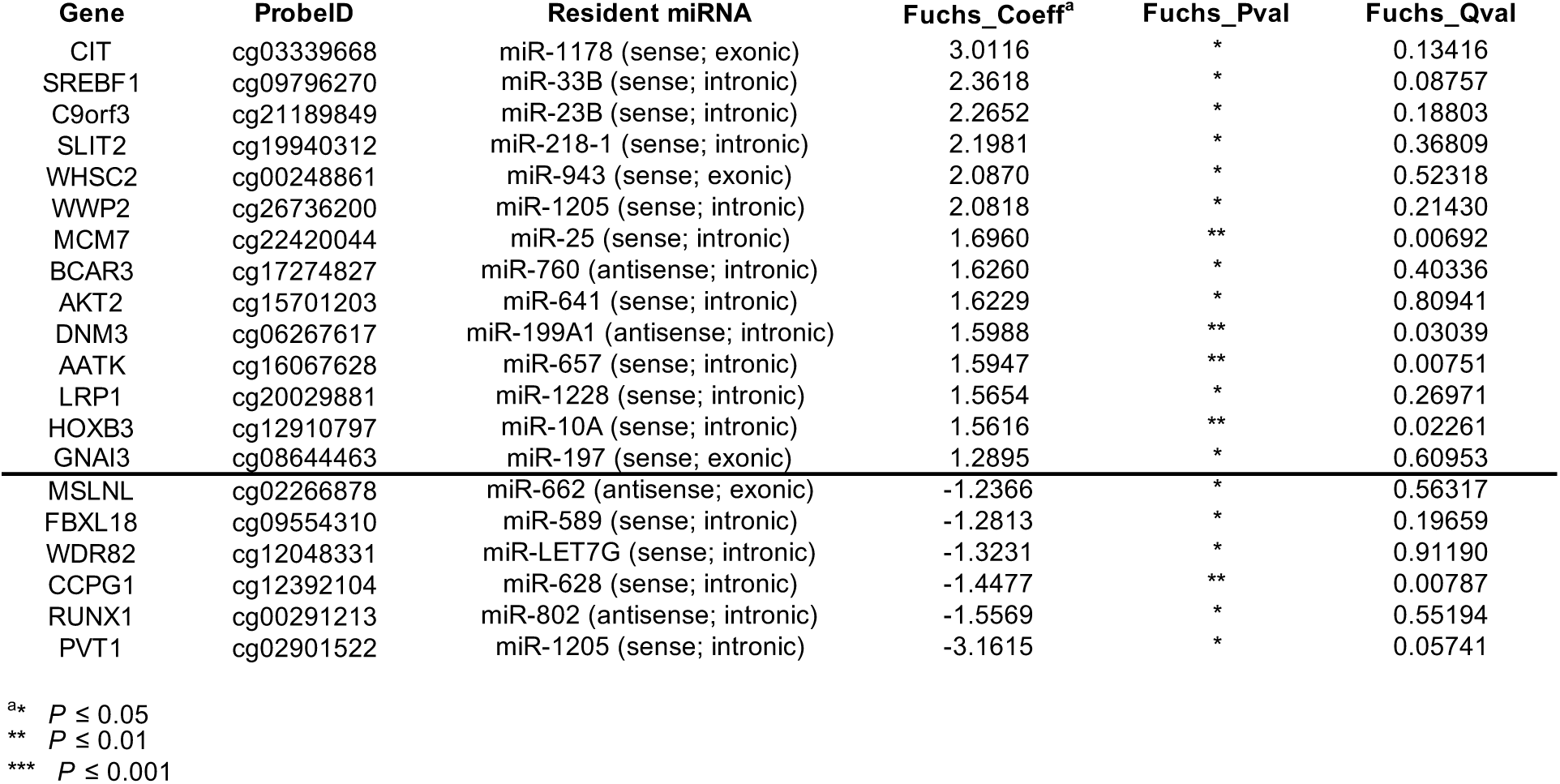
List of top 20 methylated miRNA host genes in FECD, with 14 of them being hypermethylated and 6 being hypomethylated.

### Validation of miRNA DNA methylation changes using MethyLight

To validate and quantify miRNA DNA hypermethylation changes identified by the global array [30], MethyLight analysis was performed on an additional cohort of control and FECD patient corneal samples (Table 3) [42]. MethyLight assessed the promoter DNA methylation status of following miRNAs: *miR-199A1, miR-874, miR-140, miR-23B, and miR-1306* (Fig S1, Table S1). These miRNAs were selected for validation because of their important roles in the pathogenesis of FECD or in the regulation of key cellular processes (e.g. cell survival, oxidative stress, inflammation, fibrosis, and deposition of extracellular matrix) [43–50]. The MethyLight results confirmed DNA hypermethylation in the FECD samples compared with control samples (Fig 4) and verified our array findings. All five MethyLight assays gave higher mean DNA methylation values (Percent of Methylated Reference, PMR) in the FECD samples compared to the control samples (Fig 4). *MiR-199A1* and *miR-23B* were found to be methylated in the FECD samples but unmethylated in the control samples (*P* < 0.05). For *miR-199A1*, the average PMR values were 20 and 3 for FECD and control samples, respectively (*p =* 0.039; Fig 4). For *miR-23B*, the mean PMR values were 34 for FECD samples and 5 for control samples (*p =* 0.038; Fig 4). Taken together, these MethyLight results confirmed hypermethylation of miRNA sequences in FECD tissue compared with normal control samples found in the genome-wide array using an independent set of patient samples.

**Fig 4.**
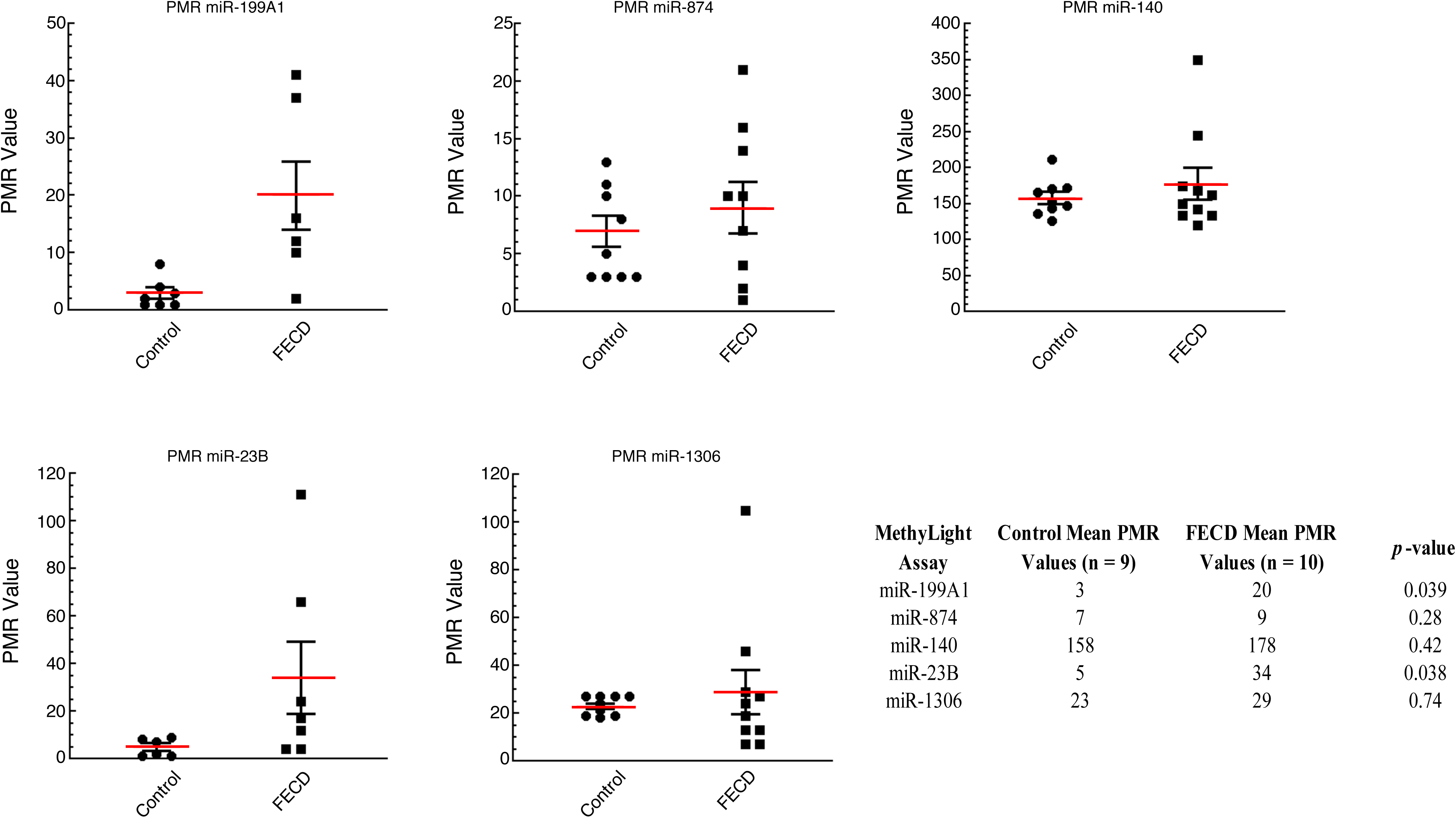
MethyLight analysis for *miR-199A1*, *miR-874*, *miR-140*, *miR-23B* and *miR-1306* in FECD and control endothelial tissues. The methylation levels of five miRNA genes were quantified by real-time PCR-based MethyLight assays on samples from control (n = 9) and FECD (n = 10) samples. MethyLight data are presented as percent of methylated reference (PMR). The table lists the mean PMR values for the control and FECD samples and p-values for each MethyLight assay.

**Table 3.** Demographics of study participants for MethyLight analysis.

|  | **All FECD (n = 12)** | **Control (n = 9)** | **Analyzed FECD (n = 10)** |
| --- | --- | --- | --- |
| **Age (SD)** | 72.12 (8.19) | 56.11 (9.05) | 73.07 (6.36) |
| Age Range | 54.7-85.1 | 36-67 | 66.1-85.1 |
| **Sex** |  |  |  |
| Male | 7 (58%) | 2 (22%) | 6 (60%) |
| Female | 5 (42%) | 7 (78%) | 4 (40%) |
| **Study Eye** |  |  |  |
| Right | 3 (25%) | 6 (67%) | 2 (29%) |
| Left | 9 (75%) | 3 (33%) | 8 (71%) |
| **Procedure** |  |  |  |
| DSAEK | 2 (17%) | N/A | 1 (10%) |
| DMEK | 10 (83%) | N/A | 9 (90%) |
| **FECD Measures** |  |  |  |
| Guttae (SD) | 2.7 (0.78) | 0 | 2.8 (0.82) |
| Pachymetry (SD) | 632 (126) | N/A | 626 (135) |

### Relationship between miRNA DNA methylation status and expression

The majority of the miRNA DNA methylation changes observed in FECD tissues occurred in miRNA promoter sequences. Because promoter DNA methylation is often inversely correlated with gene expression levels [51, 52], we next sought to determine how DNA methylation might affect miRNA expression in FECD tissue. Matthaei et al. previously compared miRNA expression profiles of corneal endothelial samples obtained from FECD patients and from normal donors using transcriptome analysis [36]. Their results demonstrated downregulation of 87 miRNAs in FECD compared with normal endothelium and suggested that altered miRNA expression may play an important role in the pathogenesis of FECD disease [36]. Therefore, we integrated our DNA methylation data with their miRNA expression data and generated a Venn diagram showing all differentially methylated and differentially expressed miRNAs (Fig 5A). Of 156 miRNAs that are hypermethylated and 87 miRNAs that have down-regulated expression in FECD, 18 miRNAs have concurrent hypermethylation and decreased expression in FECD compared to the control samples (Fig 5A, B). In particular, miR-199b-5p expression was almost completely silenced [36] and it was the miRNA with the highest level of promoter hypermethylation (Fig 5B). This strong correlation between down-regulated miR-199b-5p expression and its high level of promoter hypermethylation in FECD suggests that miR-199b-5p directed pathways may have an important role in FECD pathogenesis.

**Fig 5.**
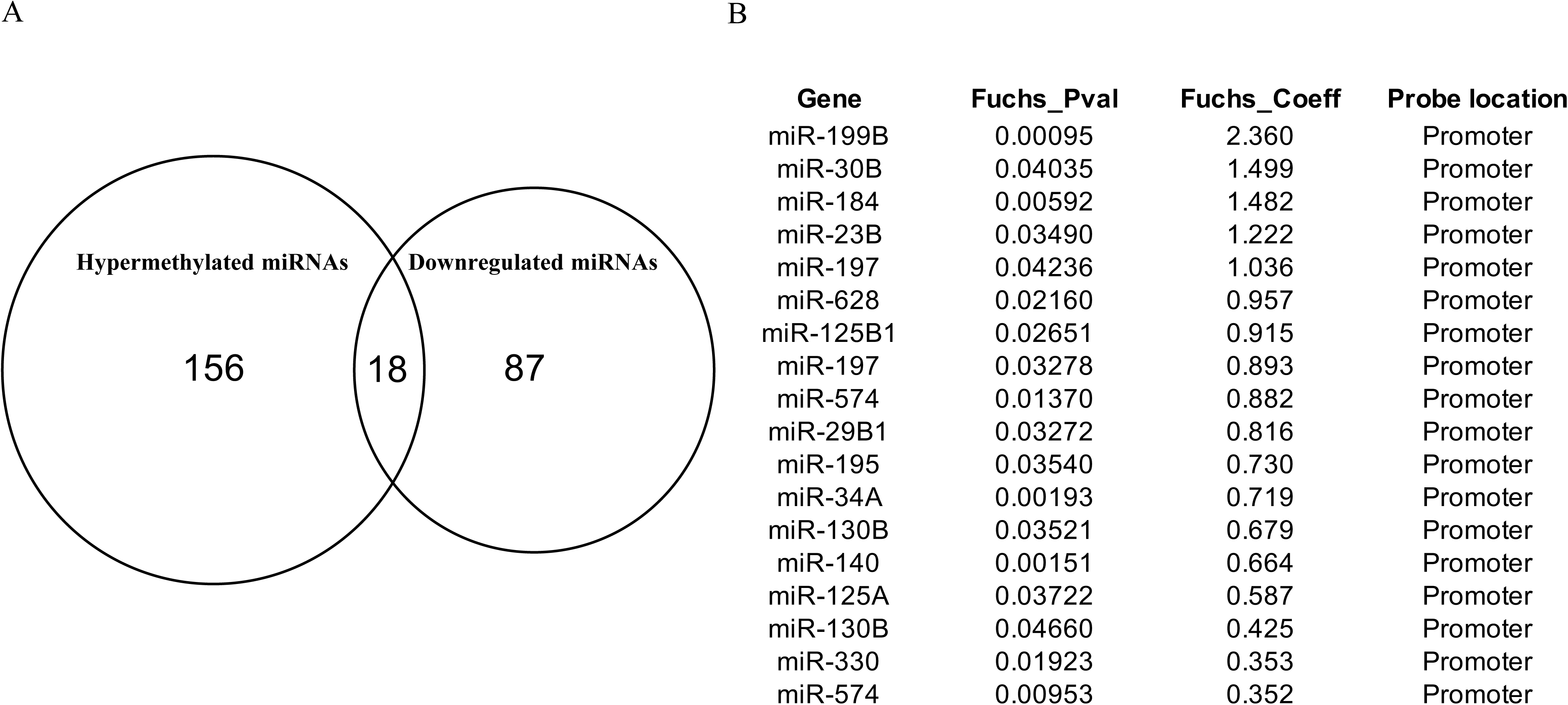
Differentially methylated and expressed miRNAs between FECD and donor samples. **(A)** Venn diagram showing the overlap of differentially methylated miRNA genes (n = 156) and differentially expressed miRNA genes (n = 87) between FECD and control samples. **(B)** List of the 18 miRNAs that have concurrent hypermethylation and decreased expression in FECD compared to the control samples.

### miR-199b-5p negatively regulates Snai1 and ZEB1 expression in corneal endothelial cells

MiRNAs can negatively regulate gene expression by directly binding to specific sequences in the 3′-UTR of target mRNAs and inducing mRNA cleavage or translation inhibition [53]. In mammals, there is a high-degree of Watson-Crick base-pairing between miRNA and target mRNA at nucleotides 2-7 at the 5’ end of miRNA, termed the “seed match” [54]. Mismatches in the miRNA-mRNA duplex were found to be ineffective in repressing gene expression [55]. To further delineate the functional role of miR-199b-5p in FECD pathogenesis, we performed *in silico* analysis to predict putative target genes and corresponding binding sites using two computational prediction algorithms (Targetscan and miRmap). More than one thousand targets of miR-199b-5p were predicted from these programs. Snai1 and ZEB1 were of particular interest because their overexpression leads to excessive extracellular matrix production in FECD [56]. Both prediction tools independently gave Snai1 and ZEB1 high scores (97 and 83.4 respectively). Sequence alignment analyses revealed a highly conserved miRNA-199b-5p binding motif in the 3′-UTR of both Snai1 and ZEB1 across many species (Fig 6A). In particular, this predicted binding site was located in the 3′-UTR of human Snai1 (positions 725-731; NM_005985.3) and ZEB1 (positions 1023-1029; NM_001128128.2) (Fig 6B).

**Fig 6.**
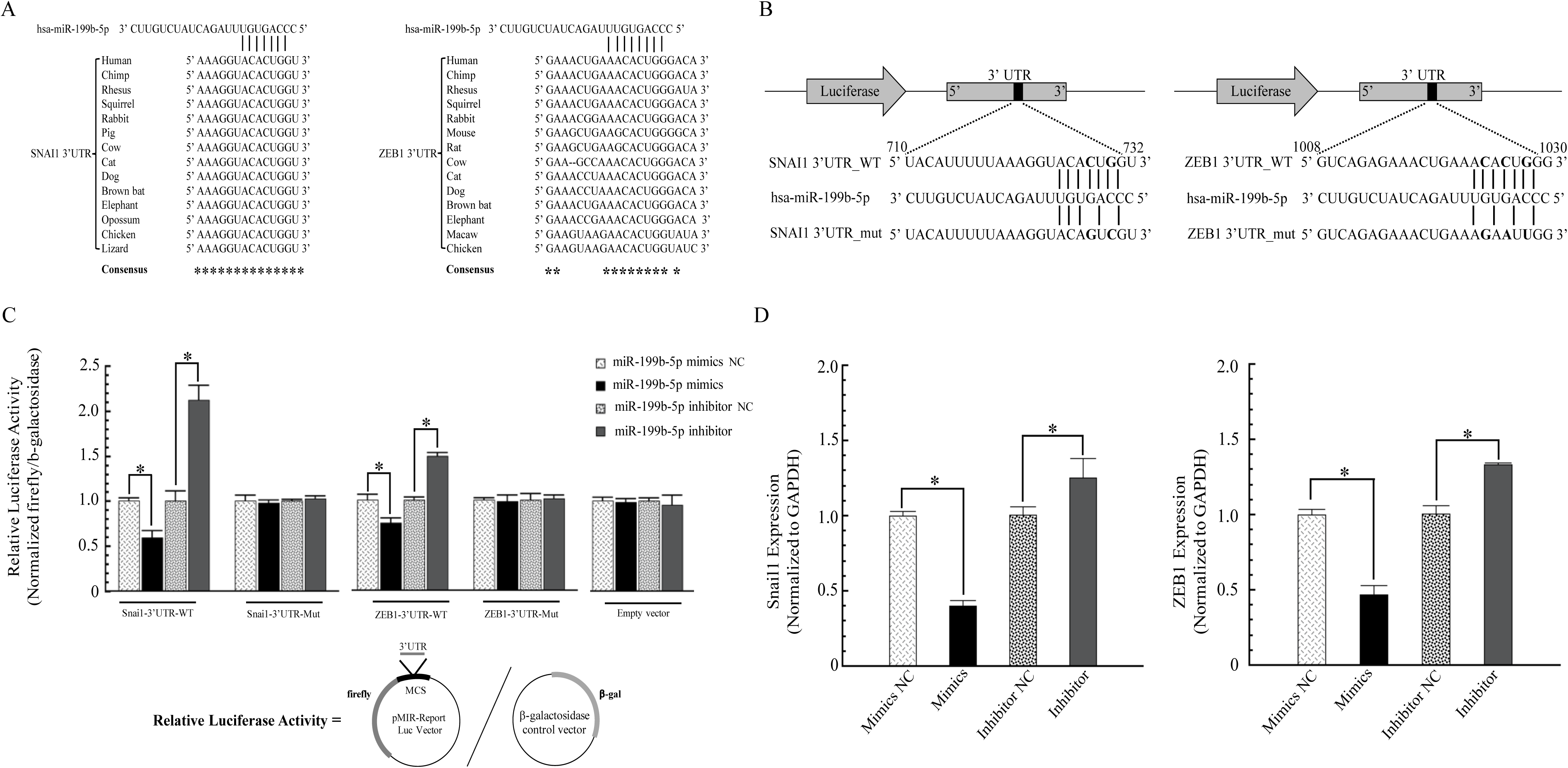
miR-199b-5p binds directly to the 3′-UTR of Snai1 and ZEB1 mRNAs. **(A)** *In silico* analyses predicted a highly conserved binding site for miR-199b-5p in the 3′-UTR of Snai1 and ZEB1 mRNAs. **(B)** The luciferase reporter plasmids containing the wild-type 3′-UTR or mutant 3′-UTR of Snail and ZEB1 with the putative binding sites for miR-199b-5p. **(C)** The direct binding of miR-199b-5p to 3′-UTR of Snai1 and ZEB1 mRNAs was detected by a dual luciferase activity assay. The luciferase reporter plasmids and β-galactosidase expression plasmids were co-transfected into HEK293 or HCEnC-21T cells with either miR-199b-5p mimic, miR-199b-5p mimic negative control (NC), miR-199b-5p inhibitor, and miR-199b-5p inhibitor negative control (NC). Luciferase activity was measured by the dual luciferase reporter assay system. Relative luciferase activities were calculated by normalizing firefly luciferase activity to β-galactosidase activity in the same sample to correct for transfection efficiencies. Data are represented as the mean ± SEM (n = 3; *P* < 0.05). **(D)** The effect of miR-199b-5p on the expression of Snail and ZEB1. HCEnC-21T cells were transfected with miR-199b-5p mimic or miR-199b-5p inhibitor with their corresponding negative controls. 48 hours post-transfection, total RNA was extracted and mRNA expression levels of Snail and ZEB1 were quantified by qRT-PCR. Data are represented as the mean ± SEM (n = 3; *P* < 0.05).

To investigate whether Snai1 and ZEB1 are direct targets of miR-199b-5p and to assess the role of miR-199b-5p in regulating Snai1 and ZEB1 expression in corneal endothelial cells, we cloned human Snai1 and ZEB1 3′-UTR sequences into a luciferase reporter vector (pMIR-Snai1-WT and pMIR-ZEB1-WT; Fig 6B, C). We additionally cloned fragments of human Snai1 and ZEB1 3′-UTR with mutated miR-199b-5p binding sites into a luciferase reporter vector (pMIR-SNAI1-Mut and pMIR-ZEB1-Mut; Fig 6B, C). The reporter plasmids (pMIR-control, pMIR-Snai1/ZEB1-WT, and pMIR-Snai1/ZEB1-Mut) were co-transfected into human corneal endothelial cells or HEK293 cells with either miR-199b-5p mimics, miR-199b-5p mimic-negative control, miR-199b-5p inhibitor, or miR-199b-5p inhibitor-negative control, along with a β-galactosidase expression plasmid as an internal control. The dual-luciferase reporter assays showed that miR-199b-5p regulated Snai1 and ZEB1 by binding directly to their 3′-UTR sequences. MiR-199b-5p mimics significantly decreased the luciferase activity approximately 50% (p = 0.039) and 30% (p = 0.009), respectively, in cells co-transfected with pMIR-Snai1-WT and pMIR-ZEB1-WT, compared with their corresponding negative controls (Fig 6C). In contrast, miR-199b-5p inhibitor significantly increased their luciferase activities by 2.1- and 1.5-fold, respectively (*P* < 0.05; Fig 6C). However, miR-199b-5p mimic and inhibitor had no effect on luciferase activities in cells co-transfected with pMIR-Snai1-Mut, pMIR-ZEB1-Mut, or empty pMIR-control vectors (*P* > 0.05; Fig 6C). Taken together, these data show that miR-199b-5p directly targets the 3′-UTRs of Snai1 and ZEB1 mRNA transcripts.

To further evaluate the effect of miR-199b-5p on Snai1 and ZEB1 expression in human corneal endothelial cells, we transfected miR-199b-5p mimic, inhibitor, or negative controls into human corneal endothelial cells and measured the expression levels of Snai1 and ZEB1 by qRT-PCR. We found that the miR-199b-5p mimic significantly inhibited Snai1 and ZEB1 expression by 50%, compared to negative control group (*P* < 0.05; Fig 6D). In contrast, the miR-199b-5p inhibitor had the opposite effect and resulted in increased Snai1 and ZEB1 expression (∼ 1.3 fold, *P* < 0.05; Fig 6D). These results demonstrate that miR-199b-5p can directly bind to and negatively regulate Snai1 and ZEB1 in human corneal endothelial cells.

## Discussion

FECD is the most common type of corneal endothelial dystrophy and a leading indication for corneal transplantation in patients in the United States [6, 57]. We previously identified global DNA methylation changes that occur in the corneal endothelial tissue of FECD patients and specifically observed a high number of DNA methylation alterations occurring in miRNA sequences [30]. This finding was intriguing because prior reports had demonstrated that miRNAs were differentially expressed in the corneal endothelium during aging [58], and that widespread downregulation of miRNA levels occurred in the corneal endothelium of patients with late-onset FECD [36, 59]. Because DNA methylation has been shown to be a mechanism for regulating miRNA expression [33], we performed a sub-analysis of the miRNA DNA methylation array data. The most differentially methylated miRNA sequences were further validated by quantitative MethyLight assay using an additional patient cohort. *MiR-199B* was identified as the most hypermethylated miRNA sequence in FECD and was selected for additional analysis because its expression was almost completely silenced in FECD [36]. *In silico* analyses identified Snai1 and ZEB1 as potential direct targets of miR-199b-5p. Using a luciferase reporter assay, we confirmed that miR-199b-5p directly targeted the 3′-UTR of both Snai1 and ZEB1 transcripts and negatively regulated their expression. Collectively, these results demonstrate that miR-199b-5p hypermethylation may contribute to late-onset FECD pathogenesis. Our findings suggest that miR-199b-5p hypermethylation leads to its down-regulated expression and consequently results in the decreased expression of miR-199b-5p target genes, including Snai1 and ZEB1.

MiRNAs are small non-coding RNAs that negatively regulate gene expression by binding to specific sequences in the 3′-UTR of target mRNAs [58, 60]. Such interactions may result in either translation inhibition or induction of mRNA cleavage [60]. Numerous studies have shown that miRNAs are evolutionarily conserved and are key regulators of diverse biological processes such as development, cell proliferation and differentiation, apoptosis and metabolism [61]. MiRNAs also have important regulatory roles in disease progression, including oncogenesis [62, 63]. The molecular mechanisms that control miRNA expression are therefore of critical importance in better understanding normal physiologic processes and disease pathogenesis. Recently, DNA methylation has emerged as a key regulatory mechanism of miRNA expression in several different tissues and disease states [62–65].

In this study, we have demonstrated aberrant DNA methylation of miRNA sequences in corneal endothelial tissue of FECD patients. Our array dataset included 2,227 probes associated with 463 miRNA genes, with multiple probes targeting single miRNA genes. We identified 216 probes associated with 156 miRNA genes that were differentially methylated between FECD and control samples, and the vast majority were hypermethylated in FECD. Furthermore, we found that the aberrant DNA methylation occurred almost exclusively in the promoter regions of miRNAs. Since promoter methylation and gene expression are usually inversely correlated, these results suggest DNA hypermethylation as a potential mechanism for the widespread downregulation of miRNA levels in FECD [36]. This preferential hypermethylation of miRNA gene promoters was also reported in other studies [33, 66-68].

To further investigate the methylated probes in a broader genomic context, we mapped the 50bp sequences of all 216 differentially methylation probes associated with 156 miRNA genes to the human genome. We found that approximately three-quarters of these probes were located within introns and/or exons of relevant host genes. The intragenic resident miRNAs on the same strand as their host genes can be co-transcribed by RNA polymerase II and co-regulated with their host genes [69]. A genome-scale DNA methylation analysis specifically on miRNA host genes revealed that miRNA host genes were frequent targets for aberrant DNA methylation and in particular downregulation of miR-10a was correlated with the promoter hypermethylation of its host gene *HOXB4* in tumorigenesis [39]. Our data found that the miRNA host genes were differentially methylated in FECD and that their gene body regions were preferential targets of aberrant methylation. Even though gene body methylation is positively correlated with gene expression [70], we were unable to measure the changes in mRNA levels of miRNA host genes on the same sample cohorts used in the DNA methylation analyses because of the low cellular yield. Therefore, the physiological relevance of DNA methylation changes of miRNA host genes in FECD remains to be further explored. Additionally, a subset of miRNA genes and their host genes shared hypermethylation of their individual promoters, suggesting that DNA methylation may play an important role in repressing the expression of certain miRNAs and their host genes simultaneously in FECD.

Using two independent DNA methylation assay technologies and two separate patient cohorts, we identified miR-199B as the most hypermethylated miRNA in the FECD samples. Interestingly, miR-199b-5p has been shown to be almost completely silenced in FECD tissues [36]. We were unable to perform side-by-side comparative miRNA transcriptome analysis on the same sample cohorts used in the DNA methylation analyses because of the low cellular yield from the FECD samples. To delineate the mechanism by which miR-199b-5p may contribute to FECD pathogenesis, we used computational algorithms to search for putative target genes, and identified Snai1 and ZEB1 as having high prediction scores. Further functional analyses using a luciferase reporter assay confirmed both 3′-UTRs of Snai1 and ZEB1 transcripts as being direct targets of miR-199b-5p. Our result is consistent with the prior finding that miR-199a-5p, a close family member of miR-199b-5p, directly binds the 3’-UTR of the Snai1 mRNA and reduces Snai1 protein level via the UGUGACC motif in its seed sequence [71]. Members of the same miRNA family can have similar physiological function and share the same predicted targets because of their conserved sequence and structural configuration [72]. Our finding that the 3’-UTRs of Snai1 and ZEB1 have the same predicted target site recognized by the identical seed sequence for both miR-199a-5p and miR-199b-5p supports the miR-199 family as having an important regulatory role in Snai1 and ZEB1 expression and function.

Snai1 and ZEB1 are zinc finger transcription factors that regulate gene expression in multiple tissues, including the cornea. Okumura et al. showed that immortalized corneal endothelial cells obtained from late-onset FECD patients highly expressed Snai1 and ZEB1 had excessive production of ECM proteins, including type I collagen and fibronectin [56]. Katikireddy et al. found that Snai1 expression level is significantly upregulated in ex vivo FECD specimens as compared to control samples [73].

A phenotypic clinical feature of FECD is the development of corneal guttae, which are abnormal collagenous excrescences of the corneal endothelial basement membrane (Descemet’s membrane). Recent studies have shown that Snail and ZEB1 can also reduce cell adhesion, increase cell migratory capacity [74–76], and promote apoptosis [77, 78]. These phenotypic features have also been observed during FECD pathogenesis [79–81].

Our findings support a model in which aberrant promoter hypermethylation of miR-199b-5p in FECD leads to the down-regulated expression of Snai1 and ZEB1 expression and consequent pathologic overproduction of ECM proteins in the cornea (Fig 7). Dysregulated DNA methylation of miRNA promoters has been found to be a biomarker in the detection, diagnosis, and prognosis of various cancers types including breast [82], gastrointestinal [83], and lung [84]. Our results provide a novel mechanistic insight into the function of DNA methylation in the pathogenesis of FECD and support further studies to determine how methylation of miR-199b-5p may be used as a clinical biomarker of phenotype expression in FECD.

**Fig 7.**
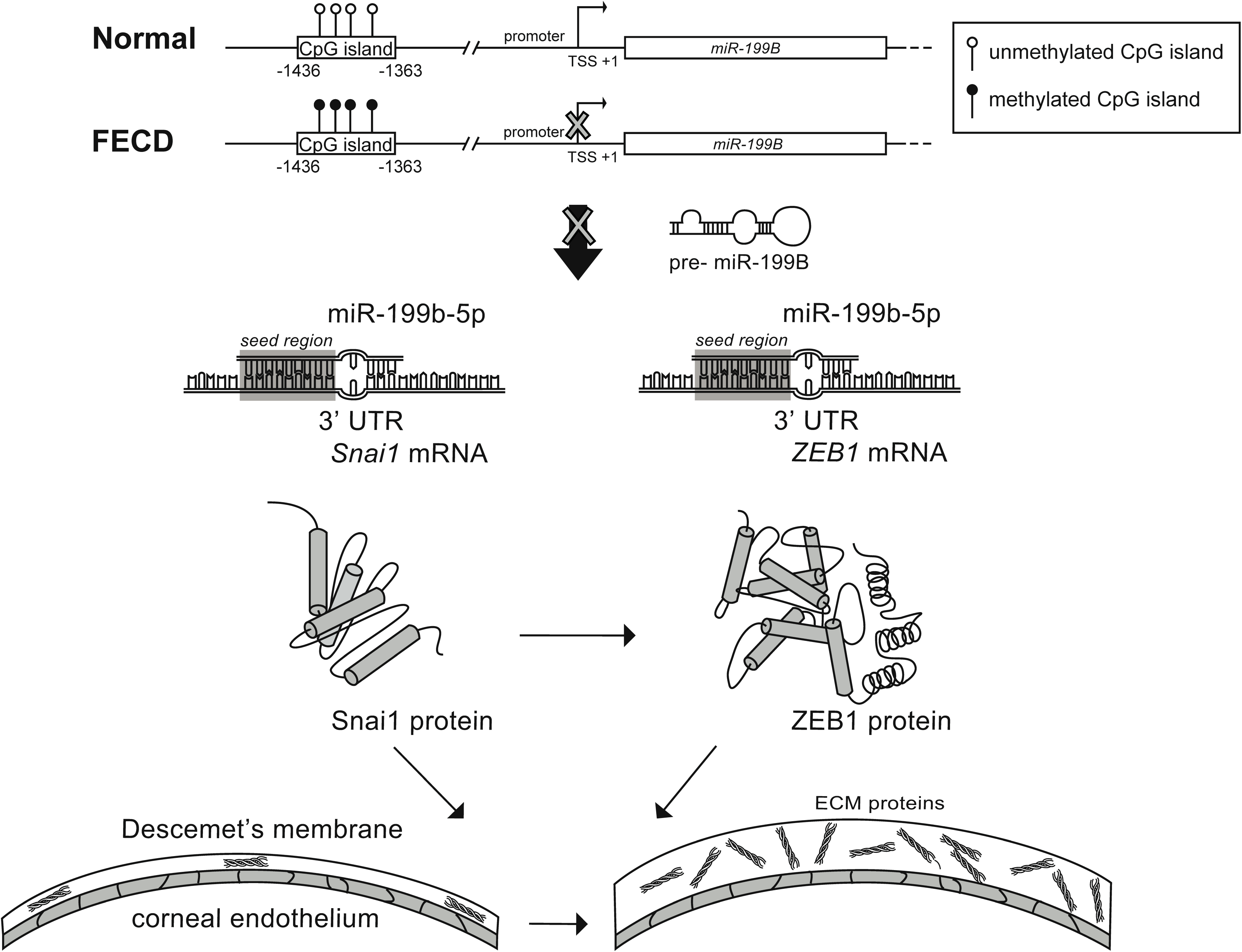
*MiR-199B* is hypermethylated in CpG island at its promoter region and its mature transcript miR-199b-5p directly inhibits the expression of Snai1 and ZEB1 in FECD. *MiR-199B* is the most hypermethylation miRNA genes in FECD. Its mature transcript miR-199b-5p functions as a direct negative regulatior of two zinc finger transcription factors, Snai1 and ZEB1, which have been shown to lead to increased production of extracellular matrix proteins in FECD.

## Materials and Methods

### Ethical compliance

Institutional Review Board (IRB)/Ethics Committee approval was obtained from the University of California, San Francisco Human Research Protection Program (Study Number 11-07020). Written informed consent was obtained from all participants. Protected health information was masked according to HIPAA privacy standards and the patient database was managed securely in Research Electronic Data Capture (REDCap) [85]. All of the described research adheres to the tenets of the Declaration of Helsinki.

### Subjects and selection criteria

Corneal endothelium was collected from FECD patients undergoing endothelial keratoplasty by two surgeons (D.G.H. and J.R.R-N.) at the University of California, San Francisco. Patients with a diagnosis of FECD and scheduled for endothelial keratoplasty between the dates of 2/12/2013 and 10/27/2014 (for Illumina Infinium HumanMethylation450 BeadChip analysis) and 1/18/2017 and 1/29/2018 (for MethyLight analysis) were recruited for the study. Written consent was obtained from participating patients and clinical information from the most recent office visit was collected from electronic medical documentation, including guttata score and pachymetry. Age- and gender-matched non-FECD corneal endothelial samples were obtained from an eye bank (SightLife, Seattle WA; and San Diego Eye Bank, San Diego CA) and processed in the same manner as the FECD samples.

### DNA methylation microarray

Array-based DNA methylation data was collected in our prior studies [30]. The IDAT files are available on the GEO DataSets database [accession number GSE94462; National Center for Biotechnology Information (NCBI), Bethesda, MD, USA].

### MethyLight assay

DNA methylation levels were measured using MethyLight technology, which is a quantitative, TaqMan-based real-time PCR assay using bisulfite converted DNA as a template [86]. Genomic DNA (200-500 ng) for each sample was converted with bisulfite using the Zymo EZ DNA Methylation kit (Zymo Research, Irvine, CA) as per the manufacturer’ instructions. M. SssI-treated DNA sample was included as a methylated reference. An interspersed *ALU* repeats-based methylation-independent reaction was also included as a normalization control. The percent of methylated reference (PMR) values for each sample were calculated for each sample as follows: PMR = 100 X (GENE/ALU)_sample_ / (GENE/ALU)_M. SssI-Reference_. The following probes targeting miRNA promoter sequences were chosen for validation with the MethyLight assay because they were significantly differentially methylated in FECD samples compared to the control samples: miR-199A1 (cg18544365), miR-874 (cg18251187), miR-140 (cg07281938), miR-23B promoter (cg00351472), and miR-1306 (cg20689730). A complete list of primers and probes for all MethyLight reactions is provided in **Supplementary Table S1**.

### Construction of the luciferase reporter plasmids

The putative binding sites in the 3′-UTRs of *Snai1* and *ZEB1* genes were bioinformatically predicted for miR-199b-5p using multiple computational prediction algorithms, including TargetScan and miRmap. The 3′-UTR sequences of both genes were then amplified from genomic DNA obtained from HEK293 cells with Phusion® High-Fidelity DNA Polymerases (NEB, Ipswich, MA) and cloned into the multiple cloning site of the pMIR-REPORT luciferase miRNA expression vector (Thermo Fisher Scientific, Waltham, MA) using the In-fusion HD Cloning Kit (Clontech, Mountain View, CA). The primers used to amplify the *Snail* 3′-UTR were: 5′-GAGTGATGAAAGCTGCGCACTAGTGGCAATTTAACAATGTCTGAAAAGG-3′, and 5′-AAAGATCCTTTATTAAGCTTCTGTACATATAACTATACAAAACGTTTCC-3′; the primers used to amplify the *ZEB1* 3′-UTR were: 5′-GAGTGATGAAAGCTGCGCACTAGTGCAGGGACTAACAATGTTAATCTG-3′, and 5′-AAAGATCCTTTATTAAGCTTCTACAGTCCAAGGCAAGTATAAATG-3′. Two mutant Snai1 and ZEB1 3′-UTR reporter vectors that lacked the binding sites for miR-199b-5p were generated using standard PCR-based overlap-extension protocols. The primers used to amplify the mutated *Snai1*3′-UTR were: *Snai1* 3′-UTR-M1, 5′-GAGTGATGAAA GCTGCGCACTAGTGGCAATTTAACAATGTCTGAAAAGG-3′, and 5′-AATACGACTGTACCTTTAAAAATGTAAAC-3′; *Snai1* 3′-UTR-M2, 5′-AAGGTACAGTCG TATTTATATTTCAAAC-3′, and 5′-AAAGATCCTTTATTAAGCTTCTGTACATATAACTATACAAAACGTTTCC-3′. The primers used to amplify the mutated *ZEB1* 3′-UTR were: *ZEB1* 3′-UTR-M1, 5′-GAGTGATGAAAGCTGCGCACTAGTGCAGGGACTAACAATGTTAATCTG-3′, and 5′-AT GTCCAATTCTTTCAGTTTCTCTGACAGAGTCAGT-3′; *ZEB1* 3′-UTR-M2, 5′-ACTGAAAGAATTGGACATTTCATCCTTCAATTCCTCGG-3′, and 5′-AAAGATCCTTTATTAAGCTTCTACAGTCCAAGGCAAGTATAAATG-3′. All clones were verified by DNA sequencing (Elim Biopharmaceuticals, Hayward, CA).

### Dual luciferase reporter assay

Dual luciferase reporter assays were performed to validate *Snail1* and *ZEB1* as *bona fide* miRNA target genes. Briefly, 0.3 × 10^6^ of HCEnC-21T cells [87] or HEK293 cells were seeded in 24-well plates and then co-transfected with 500 ng of pMIR-REPORT wild-type or mutant plasmid, 100 ng of β-gal plasmid, and 25 nmol miR-199b-5p mimic, 25 nmol scrambled mimic negative control, 50 nmol miR-199b-5p inhibitor, or 50 nmol scrambled inhibitor negative control (Life Technologies, Carlsbad, CA), using Lipofectamine 3000 (Life Technologies, Carlsbad, CA) in OptiMEM (Gibco, CA). A β-galactosidase expression plasmid was used as an internal control for transfection efficiency. Forty-eight hours after transfection, cells were subjected to lysis and firefly luciferase and β-galactosidase enzymatic activities were measured consecutively using a dual-luciferase reporter assay system (Applied Biosystems, Bedford, MA) as per the manufacturer’s instructions. Relative firefly luciferase activity (firefly luciferase activity/β-galactosidase enzymatic activities) were expressed as changes relative to that value of the negative control, which was set as 1. Three independent experiments were performed in triplicate.

### Quantitative real-time PCR (qRT-PCR)

HCEnC-21T cells were seeded in 24-well plates (0.3 × 10^6^ cells/well) and then transfected with 50 nmol miR-199-5p mimic, scrambled mimic negative control, miR-199b-5p inhibitor or scrambled inhibitor negative control using Lipofectamine RNAiMAX (Life Technologies, Carlsbad, CA). Forty-eight hours after transfection, cells were lysed and total RNA was extracted using PureLink^TM^ RNA mini Kit (Ambion, Foster City, CA). RNA concentration was measured on a NanoDrop Spectrophotometer (ThermoFisher Scientific, Wilmington, DE). Superscript III reverse transcriptase (Invitrogen) was used to generate single-stranded cDNA from 0.5 µg of total RNA with oligo dT, as per the manufacturer’s instructions. qRT-PCR was run with SYBR green PCR master mix (ThermoFisher Scientific, Wilmington, DE) using ABI Prism 7000 Real-Time PCR Detection System (Applied Biosystems, Bedford, MA). mRNA transcript abundances were determined using specific primers as follows: 1) Snai1: 5′-GACCCACACTGGCGAGAAGC-3′ and 5′-GCCTGGCACTGGTACTTCTTGACATC-3′; 2) ZEB1: 5′-GCTGGGAGGATGACACAGGAAAGG-3′ and 5′-GGTCCTCTTCAGGTGCCTCAGG-3′; 3) GAPDH: 5′-CCATCTTCCAGGAGCGAGATCCCTC-3′ and 5′-CTGCAAATGAGCCCCAGCCTTC-3′. All samples were run in triplicate. All qRT-PCR reactions were run as follows: 2 min at 50°C, 10 min at 95°C, 15 s at 95°C, and 1 min at 60°C (40 cycles) with a mixture containing 1 µl of cDNA template, 7.5 µl qPCR master mix and 266.7 nmol l^−1^ of each primer in a total volume of 15 µl (ThermoFisher Scientific, Wilmington, DE), as per the manufacturer’s instructions. Data were collected during the 1 min - 60°C extension step. Melt curves were performed using the following program: 15 s at 95°C, 2 min at 60°C, and 15 s at 95°C with a step of 0.5°C every cycle. Melting curve analyses showed no primer-dimers or non-specific products. Data are presented as fold change in gene expression normalized to GAPDH. Relative quantification of expression was calculated with the 2^-ΔΔCt^ method [88].

### Statistical analyses

Statistical analyses on genome-wide methylation data of miRNA genes were executed in R [30]. All other analyses were performed using two-tailed t-tests to compare mean values using R software. The results were statistically analyzed using Student’s t-test and p-values less than 0.05 were considered statistically significant.

## Acknowledgments

We thank Emily Khuc and Selene M. Clay for collecting patients samples for the original genome-wide DNA methylation assay.

## Author Contributions

**Conceptualization:** Peipei Pan, Matilda F. Chan.

**Data curation:** Peipei Pan, Daniel J. Weisenberger, Siyu Zheng, Marie Wolf, David G. Hwang, Jennifer R. Rose-Nussbaumer, Ula V. Jurkunas, Matilda F. Chan

**Formal analysis:** Peipei Pan, Daniel J. Weisenberger, Matilda F. Chan

**Funding acquisition:** Matilda F. Chan

**Methodology:** Peipei Pan, Daniel J. Weisenberger, Matilda F. Chan

**Project administration:** Matilda F. Chan

**Supervision:** Matilda F. Chan

**Writing – original draft:** Peipei Pan, Matilda F. Chan

**Writing – review & editing:** Peipei Pan, Daniel J. Weisenberger, Siyu Zheng, Marie Wolf, David G. Hwang, Jennifer R. Rose-Nussbaumer, Ula V. Jurkunas, Matilda F. Chan

---

**Fig S1.**
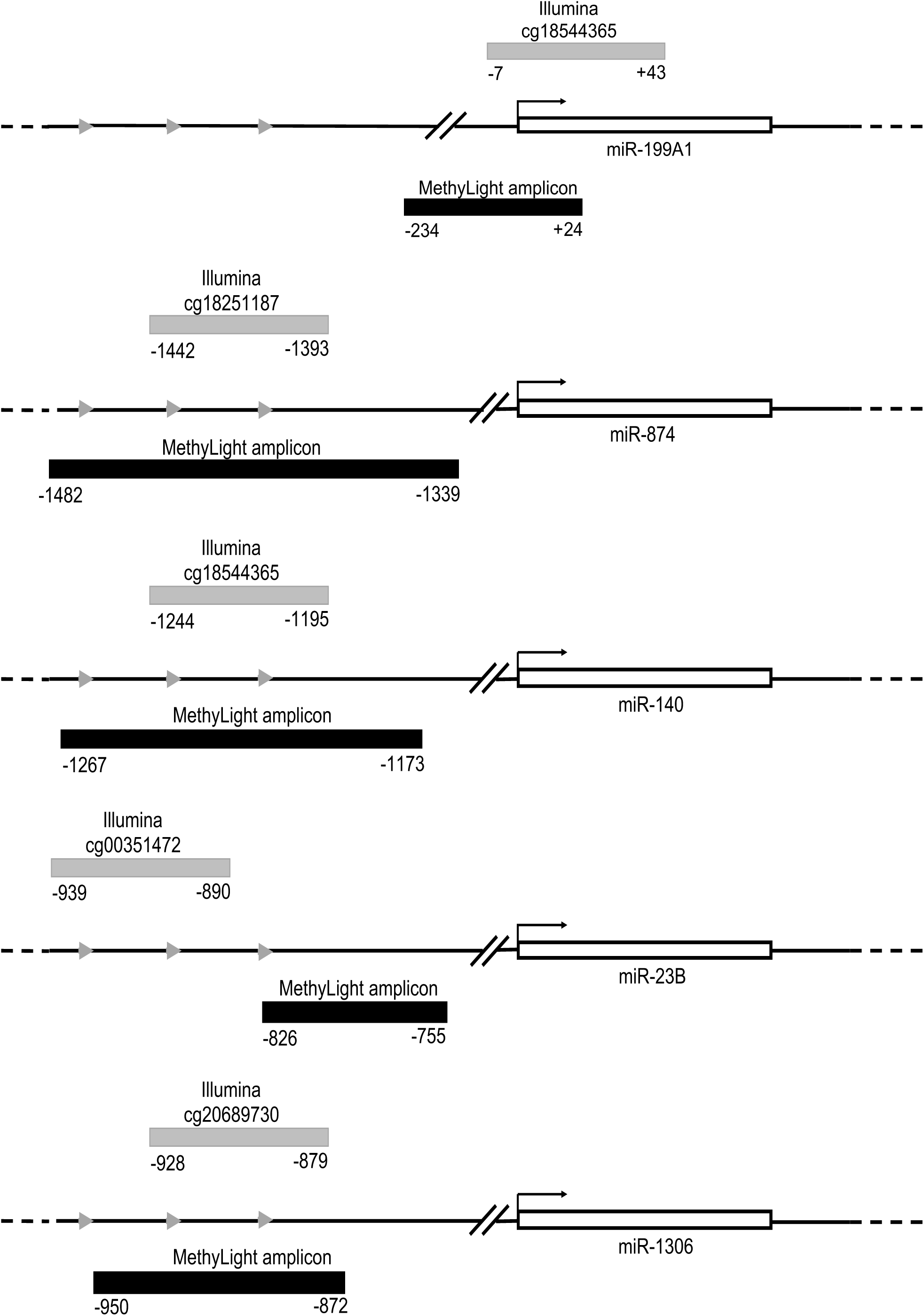
The location of Illumina Infinium HM450 array probes used in MethyLight reactions for *miR-199A1*, *miR-874*, *miR-140*, *miR-23B* and *miR-1306* genes.

**Table S1.**
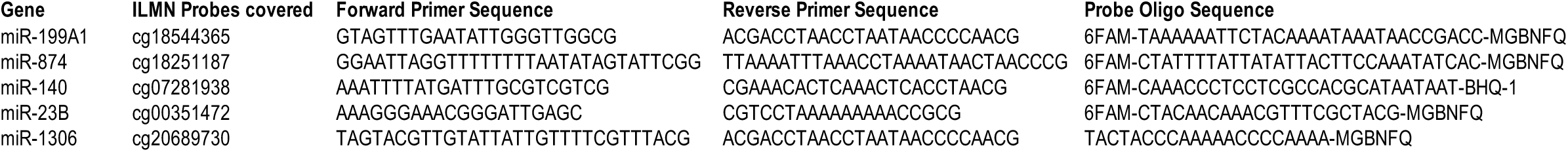
List of MethyLight primer and probe sequences.

## References

1. Bonanno JA. Molecular mechanisms underlying the corneal endothelial pump. Experimental eye research. 2012;95(1):2-7. Epub 2011/06/23. doi: 10.1016/j.exer.2011.06.004. PubMed PMID: 21693119; PubMed Central PMCID: PMCPMC3199349.

2. Bourne WM. Biology of the corneal endothelium in health and disease. Eye (Lond). 2003;17(8):912-8. doi: 10.1038/sj.eye.6700559. PubMed PMID: 14631396.

3. Adamis AP, Filatov V, Tripathi BJ, Tripathi RC. Fuchs’ endothelial dystrophy of the cornea. Survey of ophthalmology. 1993;38(2):149-68. PubMed PMID: 8235998.

4. Klintworth GK. Corneal dystrophies. Orphanet J Rare Dis. 2009;4:7. doi: 10.1186/1750-1172-4-7. PubMed PMID: 19236704; PubMed Central PMCID: PMCPMC2695576.

5. Mimura T, Yamagami S, Amano S. Corneal endothelial regeneration and tissue engineering. Progress in retinal and eye research. 2013;35:1-17. doi: 10.1016/j.preteyeres.2013.01.003. PubMed PMID: 23353595.

6. Park CY, Lee JK, Gore PK, Lim CY, Chuck RS. Keratoplasty in the United States: A 10-Year Review from 2005 through 2014. Ophthalmology. 2015;122(12):2432-42. doi: 10.1016/j.ophtha.2015.08.017. PubMed PMID: 26386848.

7. Cross HE, Maumenee AE, Cantolino SJ. Inheritance of Fuchs’ endothelial dystrophy. Arch Ophthalmol. 1971;85(3):268-72. PubMed PMID: 5313141.

8. Eghrari AO, Gottsch JD. Fuchs’ corneal dystrophy. Expert Rev Ophthalmol. 2010;5(2):147-59. doi: 10.1586/eop.10.8. PubMed PMID: 20625449; PubMed Central PMCID: PMCPMC2897712.

9. Elhalis H, Azizi B, Jurkunas UV. Fuchs endothelial corneal dystrophy. Ocul Surf. 2010;8(4):173-84. PubMed PMID: 20964980; PubMed Central PMCID: PMCPMC3061348.

10. Vithana EN, Morgan P, Sundaresan P, Ebenezer ND, Tan DT, Mohamed MD, et al. Mutations in sodium-borate cotransporter SLC4A11 cause recessive congenital hereditary endothelial dystrophy (CHED2). Nature genetics. 2006;38(7):755-7. doi: 10.1038/ng1824. PubMed PMID: 16767101.

11. Vithana EN, Morgan PE, Ramprasad V, Tan DT, Yong VH, Venkataraman D, et al. SLC4A11 mutations in Fuchs endothelial corneal dystrophy. Hum Mol Genet. 2008;17(5):656-66. doi: 10.1093/hmg/ddm337. PubMed PMID: 18024964.

12. Riazuddin SA, Zaghloul NA, Al-Saif A, Davey L, Diplas BH, Meadows DN, et al. Missense mutations in TCF8 cause late-onset Fuchs corneal dystrophy and interact with FCD4 on chromosome 9p. Am J Hum Genet. 2010;86(1):45-53. doi: 10.1016/j.ajhg.2009.12.001. PubMed PMID: 20036349; PubMed Central PMCID: PMCPMC2801746.

13. Riazuddin SA, Parker DS, McGlumphy EJ, Oh EC, Iliff BW, Schmedt T, et al. Mutations in LOXHD1, a recessive-deafness locus, cause dominant late-onset Fuchs corneal dystrophy. Am J Hum Genet. 2012;90(3):533-9. doi: 10.1016/j.ajhg.2012.01.013. PubMed PMID: 22341973; PubMed Central PMCID: PMCPMC3309196.

14. Riazuddin SA, Vasanth S, Katsanis N, Gottsch JD. Mutations in AGBL1 Cause Dominant Late-Onset Fuchs Corneal Dystrophy and Alter Protein-Protein Interaction with TCF4. American Journal of Human Genetics. 2013;93(4):758-64. doi: 10.1016/j.ajhg.2013.08.010. PubMed PMID: WOS:000326305600018.

15. Afshari NA, Igo RP, Jr., Morris NJ, Stambolian D, Sharma S, Pulagam VL, et al. Genome-wide association study identifies three novel loci in Fuchs endothelial corneal dystrophy. Nat Commun. 2017;8:14898. doi: 10.1038/ncomms14898. PubMed PMID: 28358029; PubMed Central PMCID: PMCPMC5379100.

16. Wieben ED, Aleff RA, Tosakulwong N, Butz ML, Highsmith WE, Edwards AO, et al. A Common Trinucleotide Repeat Expansion within the Transcription Factor 4 (TCF4, E2-2) Gene Predicts Fuchs Corneal Dystrophy. PLoS ONE. 2012;7(11). doi: ARTN e49083 10.1371/journal.pone.0049083. PubMed PMID: WOS:000311821000028.

17. Jurkunas UV, Rawe I, Bitar MS, Zhu C, Harris DL, Colby K, et al. Decreased expression of peroxiredoxins in Fuchs’ endothelial dystrophy. Investigative ophthalmology & visual science. 2008;49(7):2956-63. doi: 10.1167/iovs.07-1529. PubMed PMID: 18378575; PubMed Central PMCID: PMCPMC2773676.

18. Jurkunas UV, Bitar MS, Funaki T, Azizi B. Evidence of oxidative stress in the pathogenesis of fuchs endothelial corneal dystrophy. Am J Pathol. 2010;177(5):2278-89. doi: 10.2353/ajpath.2010.100279. PubMed PMID: 20847286; PubMed Central PMCID: PMCPMC2966787.

19. Buddi R, Lin B, Atilano SR, Zorapapel NC, Kenney MC, Brown DJ. Evidence of oxidative stress in human corneal diseases. J Histochem Cytochem. 2002;50(3):341-51. doi: 10.1177/002215540205000306. PubMed PMID: 11850437.

20. Wang Z, Handa JT, Green WR, Stark WJ, Weinberg RS, Jun AS. Advanced glycation end products and receptors in Fuchs’ dystrophy corneas undergoing Descemet’s stripping with endothelial keratoplasty. Ophthalmology. 2007;114(8):1453-60. doi: 10.1016/j.ophtha.2006.10.049. PubMed PMID: 17320180.

21. Feinberg AP. Phenotypic plasticity and the epigenetics of human disease. Nature. 2007;447(7143):433-40. doi: 10.1038/nature05919. PubMed PMID: 17522677.

22. Jones MJ, Goodman SJ, Kobor MS. DNA methylation and healthy human aging. Aging Cell. 2015;14(6):924-32. doi: 10.1111/acel.12349. PubMed PMID: 25913071; PubMed Central PMCID: PMCPMC4693469.

23. Pennington KL, DeAngelis MM. Epigenetic Mechanisms of the Aging Human Retina. J Exp Neurosci. 2015;9(Suppl 2):51-79. doi: 10.4137/JEN.S25513. PubMed PMID: 26966390; PubMed Central PMCID: PMCPMC4777243.

24. Welsbie DS, Yang Z, Ge Y, Mitchell KL, Zhou X, Martin SE, et al. Functional genomic screening identifies dual leucine zipper kinase as a key mediator of retinal ganglion cell death. Proc Natl Acad Sci U S A. 2013;110(10):4045-50. doi: 10.1073/pnas.1211284110. PubMed PMID: 23431148; PubMed Central PMCID: PMCPMC3593842.

25. McDonnell F, O’Brien C, Wallace D. The role of epigenetics in the fibrotic processes associated with glaucoma. J Ophthalmol. 2014;2014:750459. doi: 10.1155/2014/750459. PubMed PMID: 24800062; PubMed Central PMCID: PMCPMC3988735.

26. Baird PN, Wei L. Age-related macular degeneration and DNA methylation. Epigenomics. 2013;5(3):239-41. doi: 10.2217/epi.13.19. PubMed PMID: 23750638.

27. Gemenetzi M, Lotery AJ. The role of epigenetics in age-related macular degeneration. Eye (Lond). 2014;28(12):1407-17. doi: 10.1038/eye.2014.225. PubMed PMID: 25233816; PubMed Central PMCID: PMCPMC4268465.

28. Oliver VF, Jaffe AE, Song J, Wang G, Zhang P, Branham KE, et al. Differential DNA methylation identified in the blood and retina of AMD patients. Epigenetics. 2015;10(8):698-707. doi: 10.1080/15592294.2015.1060388. PubMed PMID: 26067391; PubMed Central PMCID: PMCPMC4622056.

29. Li F, Wang Y, Zhang G, Zhou J, Yang L, Guan H. Expression and methylation of DNA repair genes in lens epithelium cells of age-related cataract. Mutat Res. 2014;766-767:31-6. doi: 10.1016/j.mrfmmm.2014.05.010. PubMed PMID: 25847269.

30. Khuc E, Bainer R, Wolf M, Clay SM, Weisenberger DJ, Kemmer J, et al. Comprehensive characterization of DNA methylation changes in Fuchs endothelial corneal dystrophy. PLoS ONE. 2017;12(4):e0175112. doi: 10.1371/journal.pone.0175112. PubMed PMID: 28384203; PubMed Central PMCID: PMCPMC5383226.

31. Mendell JT, Olson EN. MicroRNAs in stress signaling and human disease. Cell. 2012;148(6):1172-87. doi: 10.1016/j.cell.2012.02.005. PubMed PMID: 22424228; PubMed Central PMCID: PMCPMC3308137.

32. Leung AK, Sharp PA. MicroRNA functions in stress responses. Mol Cell. 2010;40(2):205-15. doi: 10.1016/j.molcel.2010.09.027. PubMed PMID: 20965416; PubMed Central PMCID: PMCPMC2996264.

33. Lujambio A, Calin GA, Villanueva A, Ropero S, Sanchez-Cespedes M, Blanco D, et al. A microRNA DNA methylation signature for human cancer metastasis. Proceedings of the National Academy of Sciences of the United States of America. 2008;105(36):13556-61. Epub 2008/09/05. doi: 10.1073/pnas.0803055105. PubMed PMID: 18768788; PubMed Central PMCID: PMCPMC2528872.

34. Sato F, Tsuchiya S, Meltzer SJ, Shimizu K. MicroRNAs and epigenetics. FEBS J. 2011;278(10):1598-609. doi: 10.1111/j.1742-4658.2011.08089.x. PubMed PMID: 21395977.

35. Croce CM. Causes and consequences of microRNA dysregulation in cancer. Nat Rev Genet. 2009;10(10):704-14. doi: 10.1038/nrg2634. PubMed PMID: 19763153; PubMed Central PMCID: PMCPMC3467096.

36. Matthaei M, Hu J, Kallay L, Eberhart CG, Cursiefen C, Qian J, et al. Endothelial cell microRNA expression in human late-onset Fuchs’ dystrophy. Investigative ophthalmology & visual science. 2014;55(1):216-25. doi: 10.1167/iovs.13-12689. PubMed PMID: 24334445; PubMed Central PMCID: PMCPMC3888010.

37. Hinske LC, Franca GS, Torres HA, Ohara DT, Lopes-Ramos CM, Heyn J, et al. miRIAD-integrating microRNA inter- and intragenic data. Database (Oxford). 2014;2014. doi: 10.1093/database/bau099. PubMed PMID: 25288656; PubMed Central PMCID: PMCPMC4186326.

38. Dohi O, Yasui K, Gen Y, Takada H, Endo M, Tsuji K, et al. Epigenetic silencing of miR-335 and its host gene MEST in hepatocellular carcinoma. Int J Oncol. 2013;42(2):411-8. doi: 10.3892/ijo.2012.1724. PubMed PMID: 23229728; PubMed Central PMCID: PMCPMC3583616.

39. Shen J, Wang S, Zhang YJ, Kappil MA, Chen Wu H, Kibriya MG, et al. Genome-wide aberrant DNA methylation of microRNA host genes in hepatocellular carcinoma. Epigenetics. 2012;7(11):1230-7. doi: 10.4161/epi.22140. PubMed PMID: 22976466; PubMed Central PMCID: PMCPMC3499324.

40. Lutter D, Marr C, Krumsiek J, Lang EW, Theis FJ. Intronic microRNAs support their host genes by mediating synergistic and antagonistic regulatory effects. BMC Genomics. 2010;11:224. doi: 10.1186/1471-2164-11-224. PubMed PMID: 20370903; PubMed Central PMCID: PMCPMC2865499.

41. Kos A, Olde Loohuis NF, Wieczorek ML, Glennon JC, Martens GJ, Kolk SM, et al. A potential regulatory role for intronic microRNA-338-3p for its host gene encoding apoptosis-associated tyrosine kinase. PLoS ONE. 2012;7(2):e31022. doi: 10.1371/journal.pone.0031022. PubMed PMID: 22363537; PubMed Central PMCID: PMCPMC3281898.

42. Weisenberger DJ, Campan M, Long TI, Kim M, Woods C, Fiala E, et al. Analysis of repetitive element DNA methylation by MethyLight. Nucleic Acids Res. 2005;33(21):6823-36. doi: 10.1093/nar/gki987. PubMed PMID: 16326863; PubMed Central PMCID: PMCPMC1301596.

43. Lino Cardenas CL, Henaoui IS, Courcot E, Roderburg C, Cauffiez C, Aubert S, et al. miR-199a-5p Is upregulated during fibrogenic response to tissue injury and mediates TGFbeta-induced lung fibroblast activation by targeting caveolin-1. PLoS Genet. 2013;9(2):e1003291. doi: 10.1371/journal.pgen.1003291. PubMed PMID: 23459460; PubMed Central PMCID: PMCPMC3573122 in Idiopathic Pulmonary Fibrosis.

44. Shatseva T, Lee DY, Deng Z, Yang BB. MicroRNA miR-199a-3p regulates cell proliferation and survival by targeting caveolin-2. Journal of cell science. 2011;124(Pt 16):2826-36. doi: 10.1242/jcs.077529. PubMed PMID: 21807947.

45. Zhou Y, Pang B, Xiao Y, Zhou S, He B, Zhang F, et al. The protective microRNA-199a-5p-mediated unfolded protein response in hypoxic cardiomyocytes is regulated by STAT3 pathway. J Physiol Biochem. 2019;75(1):73-81. doi: 10.1007/s13105-018-0657-6. PubMed PMID: 30426460.

46. Lin N, Li XY, Zhang HM, Yang Z, Su Q. microRNA-199a-5p mediates high glucose-induced reactive oxygen species production and apoptosis in INS-1 pancreatic beta-cells by targeting SIRT1. Eur Rev Med Pharmacol Sci. 2017;21(5):1091-8. PubMed PMID: 28338182.

47. Wang L, Gao W, Hu F, Xu Z, Wang F. MicroRNA-874 inhibits cell proliferation and induces apoptosis in human breast cancer by targeting CDK9. FEBS Lett. 2014;588(24):4527-35. doi: 10.1016/j.febslet.2014.09.035. PubMed PMID: 25281924.

48. Liu QQ, Ren K, Liu SH, Li WM, Huang CJ, Yang XH. MicroRNA-140-5p aggravates hypertension and oxidative stress of atherosclerosis via targeting Nrf2 and Sirt2. Int J Mol Med. 2019;43(2):839-49. doi: 10.3892/ijmm.2018.3996. PubMed PMID: 30483753; PubMed Central PMCID: PMCPMC6317688.

49. Wang P, Zhang J, Zhang L, Zhu Z, Fan J, Chen L, et al. MicroRNA 23b regulates autophagy associated with radioresistance of pancreatic cancer cells. Gastroenterology. 2013;145(5):1133-43 e12. doi: 10.1053/j.gastro.2013.07.048. PubMed PMID: 23916944.

50. He ZJ, Li W, Chen H, Wen J, Gao YF, Liu YJ. miR-1306-3p targets FBXL5 to promote metastasis of hepatocellular carcinoma through suppressing snail degradation. Biochem Biophys Res Commun. 2018;504(4):820-6. doi: 10.1016/j.bbrc.2018.09.059. PubMed PMID: 30219228.

51. Jones PA. Functions of DNA methylation: islands, start sites, gene bodies and beyond. Nat Rev Genet. 2012;13(7):484-92. doi: 10.1038/nrg3230. PubMed PMID: 22641018.

52. Meissner A, Mikkelsen TS, Gu H, Wernig M, Hanna J, Sivachenko A, et al. Genome-scale DNA methylation maps of pluripotent and differentiated cells. Nature. 2008;454(7205):766-70. doi: 10.1038/nature07107. PubMed PMID: 18600261; PubMed Central PMCID: PMCPMC2896277.

53. Nam JW, Rissland OS, Koppstein D, Abreu-Goodger C, Jan CH, Agarwal V, et al. Global analyses of the effect of different cellular contexts on microRNA targeting. Mol Cell. 2014;53(6):1031-43. doi: 10.1016/j.molcel.2014.02.013. PubMed PMID: 24631284; PubMed Central PMCID: PMCPMC4062300.

54. Lewis BP, Burge CB, Bartel DP. Conserved seed pairing, often flanked by adenosines, indicates that thousands of human genes are microRNA targets. Cell. 2005;120(1):15-20. doi: 10.1016/j.cell.2004.12.035. PubMed PMID: 15652477.

55. Doench JG, Sharp PA. Specificity of microRNA target selection in translational repression. Genes & development. 2004;18(5):504-11. doi: 10.1101/gad.1184404. PubMed PMID: 15014042; PubMed Central PMCID: PMCPMC374233.

56. Okumura N, Minamiyama R, Ho LT, Kay EP, Kawasaki S, Tourtas T, et al. Involvement of ZEB1 and Snail1 in excessive production of extracellular matrix in Fuchs endothelial corneal dystrophy. Lab Invest. 2015;95(11):1291-304. doi: 10.1038/labinvest.2015.111. PubMed PMID: 26302187.

57. Musch DC, Niziol LM, Stein JD, Kamyar RM, Sugar A. Prevalence of corneal dystrophies in the United States: estimates from claims data. Investigative ophthalmology & visual science. 2011;52(9):6959-63. doi: 10.1167/iovs.11-7771. PubMed PMID: 21791583; PubMed Central PMCID: PMCPMC3175990.

58. Zhao X, Huang Y, Wang Y, Chen P, Yu Y, Song Z. MicroRNA profile comparison of the corneal endothelia of young and old mice: implications for senescence of the corneal endothelium. Molecular vision. 2013;19:1815-25. PubMed PMID: 23946636; PubMed Central PMCID: PMCPMC3742134.

59. Toyono T, Usui T, Villarreal G, Jr., Kallay L, Matthaei M, Vianna LM, et al. MicroRNA-29b Overexpression Decreases Extracellular Matrix mRNA and Protein Production in Human Corneal Endothelial Cells. Cornea. 2016;35(11):1466-70. doi: 10.1097/ICO.0000000000000954. PubMed PMID: 27490049; PubMed Central PMCID: PMCPMC5067961.

60. He L, Hannon GJ. MicroRNAs: small RNAs with a big role in gene regulation. Nat Rev Genet. 2004;5(7):522-31. doi: 10.1038/nrg1379. PubMed PMID: 15211354.

61. Vidigal JA, Ventura A. The biological functions of miRNAs: lessons from in vivo studies. Trends Cell Biol. 2015;25(3):137-47. doi: 10.1016/j.tcb.2014.11.004. PubMed PMID: 25484347; PubMed Central PMCID: PMCPMC4344861.

62. Asadzadeh Z, Mansoori B, Mohammadi A, Aghajani M, Haji-Asgarzadeh K, Safarzadeh E, et al. microRNAs in cancer stem cells: Biology, pathways, and therapeutic opportunities. J Cell Physiol. 2018. doi: 10.1002/jcp.27885. PubMed PMID: 30537109.

63. Chuang JC, Jones PA. Epigenetics and microRNAs. Pediatr Res. 2007;61(5 Pt 2):24R-9R. doi: 10.1203/pdr.0b013e3180457684. PubMed PMID: 17413852.

64. Gulyaeva LF, Kushlinskiy NE. Regulatory mechanisms of microRNA expression. J Transl Med. 2016;14(1):143. doi: 10.1186/s12967-016-0893-x. PubMed PMID: 27197967; PubMed Central PMCID: PMCPMC4873990.

65. Saito Y, Liang G, Egger G, Friedman JM, Chuang JC, Coetzee GA, et al. Specific activation of microRNA-127 with downregulation of the proto-oncogene BCL6 by chromatin-modifying drugs in human cancer cells. Cancer Cell. 2006;9(6):435-43. doi: 10.1016/j.ccr.2006.04.020. PubMed PMID: 16766263.

66. Gao W, Zhang C, Li W, Li H, Sang J, Zhao Q, et al. Promoter Methylation-Regulated miR-145-5p Inhibits Laryngeal Squamous Cell Carcinoma Progression by Targeting FSCN1. Mol Ther. 2019;27(2):365-79. doi: 10.1016/j.ymthe.2018.09.018. PubMed PMID: 30341010; PubMed Central PMCID: PMCPMC6369713.

67. Villela D, Ramalho RF, Silva AR, Brentani H, Suemoto CK, Pasqualucci CA, et al. Differential DNA Methylation of MicroRNA Genes in Temporal Cortex from Alzheimer’s Disease Individuals. Neural Plast. 2016;2016:2584940. doi: 10.1155/2016/2584940. PubMed PMID: 27213057; PubMed Central PMCID: PMCPMC4861808.

68. Shen J, Wang S, Siegel AB, Remotti H, Wang Q, Sirosh I, et al. Genome-Wide Expression of MicroRNAs Is Regulated by DNA Methylation in Hepatocarcinogenesis. Gastroenterol Res Pract. 2015;2015:230642. doi: 10.1155/2015/230642. PubMed PMID: 25861255; PubMed Central PMCID: PMCPMC4377534.

69. Schanen BC, Li X. Transcriptional regulation of mammalian miRNA genes. Genomics. 2011;97(1):1-6. doi: 10.1016/j.ygeno.2010.10.005. PubMed PMID: 20977933; PubMed Central PMCID: PMCPMC3019299.

70. Yang X, Han H, De Carvalho DD, Lay FD, Jones PA, Liang G. Gene body methylation can alter gene expression and is a therapeutic target in cancer. Cancer Cell. 2014;26(4):577-90. doi: 10.1016/j.ccr.2014.07.028. PubMed PMID: 25263941; PubMed Central PMCID: PMCPMC4224113.

71. Suzuki T, Mizutani K, Minami A, Nobutani K, Kurita S, Nagino M, et al. Suppression of the TGF-beta1-induced protein expression of SNAI1 and N-cadherin by miR-199a. Genes Cells. 2014;19(9):667-75. doi: 10.1111/gtc.12166. PubMed PMID: 25041364.

72. Bartel DP. MicroRNAs: target recognition and regulatory functions. Cell. 2009;136(2):215-33. doi: 10.1016/j.cell.2009.01.002. PubMed PMID: 19167326; PubMed Central PMCID: PMCPMC3794896.

73. Katikireddy KR, White TL, Miyajima T, Vasanth S, Raoof D, Chen Y, et al. NQO1 downregulation potentiates menadione-induced endothelial-mesenchymal transition during rosette formation in Fuchs endothelial corneal dystrophy. Free Radic Biol Med. 2018;116:19-30. doi: 10.1016/j.freeradbiomed.2017.12.036. PubMed PMID: 29294389; PubMed Central PMCID: PMCPMC5815941.

74. Kaufhold S, Bonavida B. Central role of Snail1 in the regulation of EMT and resistance in cancer: a target for therapeutic intervention. J Exp Clin Cancer Res. 2014;33:62. doi: 10.1186/s13046-014-0062-0. PubMed PMID: 25084828; PubMed Central PMCID: PMCPMC4237825.

75. Drake JM, Strohbehn G, Bair TB, Moreland JG, Henry MD. ZEB1 enhances transendothelial migration and represses the epithelial phenotype of prostate cancer cells. Mol Biol Cell. 2009;20(8):2207-17. doi: 10.1091/mbc.E08-10-1076. PubMed PMID: 19225155; PubMed Central PMCID: PMCPMC2669028.

76. Mathow D, Chessa F, Rabionet M, Kaden S, Jennemann R, Sandhoff R, et al. Zeb1 affects epithelial cell adhesion by diverting glycosphingolipid metabolism. EMBO Rep. 2015;16(3):321-31. doi: 10.15252/embr.201439333. PubMed PMID: 25643708; PubMed Central PMCID: PMCPMC4364871.

77. Gu Y, Zhao Y, Zhou Y, Xie Y, Ju P, Long Y, et al. Zeb1 Is a Potential Regulator of Six2 in the Proliferation, Apoptosis and Migration of Metanephric Mesenchyme Cells. Int J Mol Sci. 2016;17(8). doi: 10.3390/ijms17081283. PubMed PMID: 27509493; PubMed Central PMCID: PMCPMC5000680.

78. David CJ, Huang YH, Chen M, Su J, Zou Y, Bardeesy N, et al. TGF-beta Tumor Suppression through a Lethal EMT. Cell. 2016;164(5):1015-30. doi: 10.1016/j.cell.2016.01.009. PubMed PMID: 26898331; PubMed Central PMCID: PMCPMC4801341.

79. Loginov VI, Burdennyy AM, Pronina IV, Khokonova VV, Kurevljov SV, Kazubskaya TP, et al. [Novel miRNA genes hypermethylated in breast cancer]. Mol Biol (Mosk). 2016;50(5):797-802. doi: 10.7868/S0026898416050104. PubMed PMID: 27830681.

80. Zhong W, Li B, Xu Y, Yang P, Chen R, Wang Z, et al. Hypermethylation of the Micro-RNA 145 Promoter Is the Key Regulator for NLRP3 Inflammasome-Induced Activation and Plaque Formation. JACC Basic Transl Sci. 2018;3(5):604-24. doi: 10.1016/j.jacbts.2018.06.004. PubMed PMID: 30456333; PubMed Central PMCID: PMCPMC6234615.

81. Pigazzi M, Manara E, Bresolin S, Tregnago C, Beghin A, Baron E, et al. MicroRNA-34b promoter hypermethylation induces CREB overexpression and contributes to myeloid transformation. Haematologica. 2013;98(4):602-10. doi: 10.3324/haematol.2012.070664. PubMed PMID: 23100280; PubMed Central PMCID: PMCPMC3659992.

82. Oltra SS, Pena-Chilet M, Vidal-Tomas V, Flower K, Martinez MT, Alonso E, et al. Methylation deregulation of miRNA promoters identifies miR124-2 as a survival biomarker in Breast Cancer in very young women. Sci Rep. 2018;8(1):14373. doi: 10.1038/s41598-018-32393-3. PubMed PMID: 30258192; PubMed Central PMCID: PMCPMC6158237.

83. Toiyama Y, Okugawa Y, Goel A. DNA methylation and microRNA biomarkers for noninvasive detection of gastric and colorectal cancer. Biochem Biophys Res Commun. 2014;455(1-2):43-57. doi: 10.1016/j.bbrc.2014.08.001. PubMed PMID: 25128828; PubMed Central PMCID: PMCPMC4250419.

84. Su Y, Fang H, Jiang F. Integrating DNA methylation and microRNA biomarkers in sputum for lung cancer detection. Clin Epigenetics. 2016;8:109. doi: 10.1186/s13148-016-0275-5. PubMed PMID: 27777637; PubMed Central PMCID: PMCPMC5070138.

85. Harris PA, Taylor R, Thielke R, Payne J, Gonzalez N, Conde JG. Research electronic data capture (REDCap)--a metadata-driven methodology and workflow process for providing translational research informatics support. J Biomed Inform. 2009;42(2):377-81. doi: 10.1016/j.jbi.2008.08.010. PubMed PMID: 18929686; PubMed Central PMCID: PMC2700030.

86. Eads CA, Danenberg KD, Kawakami K, Saltz LB, Blake C, Shibata D, et al. MethyLight: a high-throughput assay to measure DNA methylation. Nucleic Acids Res. 2000;28(8):E32. PubMed PMID: 10734209; PubMed Central PMCID: PMCPMC102836.

87. Schmedt T, Chen Y, Nguyen TT, Li S, Bonanno JA, Jurkunas UV. Telomerase immortalization of human corneal endothelial cells yields functional hexagonal monolayers. PLoS ONE. 2012;7(12):e51427. doi: 10.1371/journal.pone.0051427. PubMed PMID: 23284695; PubMed Central PMCID: PMCPMC3528758.

88. Livak KJ, Schmittgen TD. Analysis of relative gene expression data using real-time quantitative PCR and the 2(-Delta Delta C(T)) Method. Methods. 2001;25(4):402-8. doi: 10.1006/meth.2001.1262. PubMed PMID: 11846609.

